# Acoustically activatable drug-loaded nanodroplets for mechanochemical therapy in solid tumors

**DOI:** 10.64898/2026.04.20.719550

**Authors:** Tiran Bercovici, Mike Bismuth, Meir Goldsmith, Dan Peer, Tali Ilovitsh

## Abstract

Stimulus-responsive nanomedicines promise spatiotemporally controlled therapy, yet most systems rely on passive delivery and lack precise, externally programmable activation while maintaining clinical compatibility. Here we engineer sub-200 nm, perfluorocarbon (PFC)-core nanodroplets (NDs) that integrate efficient core drug loading, physiological stability, and acoustically programmable activation within a single nanoscale agent. These NDs are fabricated using microfluidic nanoassembly to achieve controlled size and composition, and are designed to encapsulate fluorinated payloads directly within the liquid core. Upon exposure to a sequential dual-frequency ultrasound (US) paradigm, the NDs undergo acoustic droplet vaporization followed by low-frequency cavitation, enabling spatially confined mechanical disruption and on-demand payload release within clinically relevant acoustic limits. These properties are engineered to overcome physicochemical barriers in solid tumors, including dense extracellular matrix and restricted drug penetration. This approach achieves enhanced payload release and induces potent mechanochemical cytotoxicity in vitro. In vivo, NDs exhibit prolonged circulation and tumor accumulation, while US activation drives substantial tissue fractionation, control drug release, and increases subsequent nanoparticle uptake. When applied to a solid tumor model, this combined mechanochemical strategy improves tumor control and significantly extends survival compared to either modality alone. These acoustically activatable NDs provide a versatile system for stimulus-responsive, site-targeted drug delivery and mechanical tumor disruption, with strong potential for clinical translation.

**Graphical abstract:** 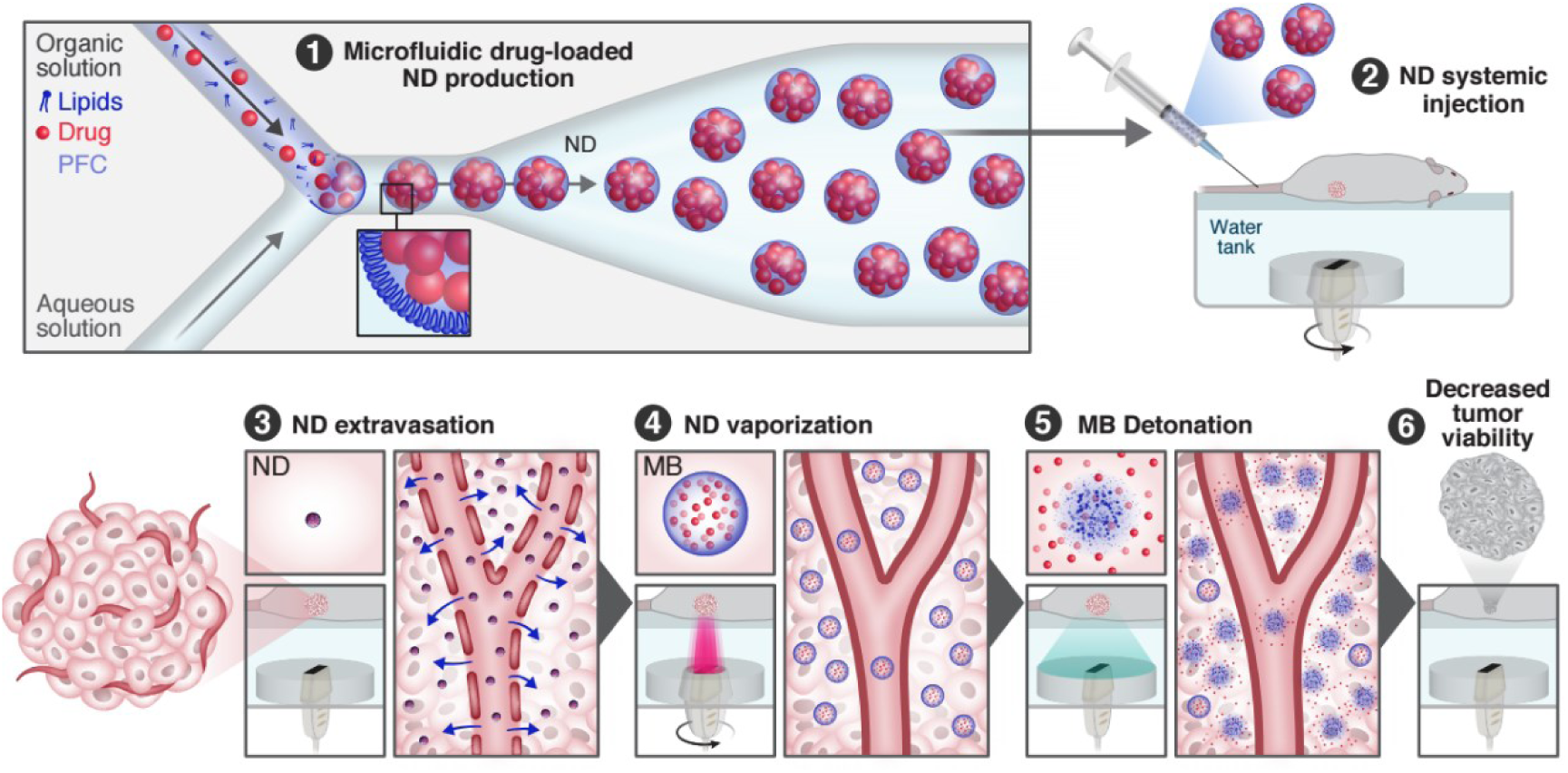

## Introduction

Solid tumors remain therapeutically challenging due to their immunosuppressive microenvironment and pronounced physicochemical barriers, including dense extracellular matrix that restrict drug penetration and reduce therapeutic efficacy^1–7^. Effective cancer treatment requires a combination of strategies to reduce tumor burden and adjuvant therapies aimed at eliminating residual malignant cells. Current cancer treatment strategies include a combination of surgery, radiotherapy, and chemotherapy, with chemotherapy being one of the most widely used approaches^8,9^. However, conventional chemotherapy often induces severe off-target toxicity due to the nonspecific systemic distribution of cytotoxic drugs^10,11^. To mitigate this limitation, nanocarrier-based drug delivery systems have been developed to improve tumor localization and reduce systemic exposure^12–15^. However, most clinically deployed nanomedicines rely on passive delivery mechanisms and lack user-controlled, externally triggered drug release.

US is the most widely used medical imaging modality due to its safety, cost effectiveness, and noninvasive nature^16^. When a US beam is focused into a confined focal region, it can induce localized bioeffects that enable therapeutic action through thermal or mechanical mechanisms^17^. These effects can be exploited to trigger site-specific uncaging of nanomedicine. Thermal approaches, such as temperature-sensitive liposomes, exploit US-induced hyperthermia to release encapsulated drugs^18^. While effective, these strategies often require prolonged sonication and are susceptible to heat diffusion, which can lead to unintended drug release in surrounding tissues^19^. Mechanical bioeffects provide an alternative activation mechanism, enabling localized cavitation and tissue disruption without bulk heating. However, existing US-responsive particles do not simultaneously achieve high drug-loading efficiency, physiological stability, and predictable acoustic activation^20,21^. Microbubbles (MB), the most commonly used US contrast agents, consist of a gas core stabilized by a shell and typically measure 1-10 μm in diameter. Under US excitation, MBs act as cavitation nuclei ^22–24^, yet they are suboptimal drug carriers, as most drug-loading strategies are confined to the shell^25^. Nanobubbles offer improved tumor penetration via the enhanced permeability and retention (EPR) effect^26^, although their drug encapsulation capacity is even lower than that of MBs, due to their smaller size^20,27^. Advanced liposomal systems enable controlled drug release but lack the mechanical bioeffects necessary for tumor debulking^18,19^.

Nanodroplets (NDs), consisting of a liquid perfluorocarbon (PFC) core stabilized by a shell, were introduced to address some of these limitations^28–30^. Upon exposure to US, NDs can undergo acoustic droplet vaporization (ADV), transitioning from a liquid droplet into a gas MB^31^. The resulting bubble can serve as US imaging contrast and enhance mechanical effects such as cavitation and tissue disruption^20,32^. Because NDs are smaller than MBs, they can circulate longer in the bloodstream, extravasate into tumors via the EPR effect, and then be activated on demand in the target tissue^30,31,33,34^. Despite these advantages, current ND platforms for drug delivery face multiple challenges. First, many formulations suffer from limited size control and high polydispersity. Most reported NDs are fabricated using bulk or emulsion-based methods such as sonication, homogenization, thin-film hydration, or MB condensation. These approaches provide limited control over nucleation and interfacial assembly, resulting in broad size distributions, batch-to-batch variability, and often increased particle diameters^35–37^. Recent work with droplet-based microfluidics has shown that it is possible to generate highly uniform droplets with tunable size and composition, opening the door to more reproducible and controllable ND formulations^36,38–40^. Second, ND vaporization behavior is strongly dependent on multiple parameters, including the PFC core, ND diameter, US frequency, and applied peak negative pressure (PNP)^41,42^. Condensation-based techniques commonly employ perfluorobutane (C_4_F_10_), which has a low boiling point (−1.7 °C) and exists as a gas at room temperature. While such formulations have demonstrated efficacy for US mechanotherapy^43^, they can be less stable under physiological conditions and prone to spontaneous vaporization^28^. To improve in vivo stability, higher boiling point PFCs such as perfluoropentane (C_5_F_12_; boiling point ∼29 °C) and perfluorohexane (C_6_F_14_; boiling point ∼56 °C) are therefore often used^28,44^. However, this increased stability comes at the cost of higher vaporization thresholds^28,44^. Third, while we have shown that insonating MBs at low frequencies enables noninvasive US surgery by significantly reducing the PNP required to induce cavitation^24,45^, ND vaporization typically requires PNPs in the MPa range^44,46–48^. At low frequencies, these pressures often exceed FDA-approved safety limits for diagnostic imaging (mechanical index < 1.9). This challenge is further amplified for smaller NDs and higher boiling point PFCs, which require even higher PNPs for vaporization^44,49^. Finally, efficient drug loading into the PFC core remains a major challenge. Most existing approaches rely on shell loading or double-emulsion strategies, which either limit payload capacity or increase droplet diameter, thereby compromising tumor penetration and systemic delivery^20,50^.

Here we introduce an acoustically activatable NDs platform designed to enable controlled droplet fabrication, efficient drug loading into the core, and enhanced US-triggered therapeutic activation within a single nanoscale agent. We employ microfluidic nanoassembly to achieve precise control over ND size (<200 nm), composition, and concentration, enabling reproducible vaporization behavior and enhanced tumor extravasation. To ensure physiological stability while preserving acoustic responsiveness, higher-order PFC cores are incorporated. Efficient drug loading is achieved through rational selection of fluorinated therapeutic molecules that exhibit physicochemical compatibility with the PFC phase, enabling dissolution within the droplet core without increasing ND size. Fluorination has been shown to enhance metabolic stability and pharmacokinetic performance, with many fluorinated drugs already approved or in clinical use^51–58^. Using this strategy, we develop three classes of NDs, including blank NDs, fluorophore-loaded NDs incorporating BODIPY dyes for mechanistic characterization, and chemotherapy-loaded NDs using 5-fluorouracil (5-FU) as a model cytotoxic agent. To enable treatment within FDA safety limits, we implement a dual-frequency US strategy (Figure 1). A rotating MHz-frequency imaging transducer first induces controlled ADV within FDA mechanical index constraints, vaporizing NDs into MBs. Subsequently, a low-frequency therapeutic transducer drives vaporized ND collapse, generating potent mechanical fractionation and on-demand payload release at the target site. This sequential activation paradigm reduces the required PNP for therapeutic bioeffects while preserving spatial precision and safety. Together, this integrated design establishes a generalizable framework for low-energy, mechanically active US nanomedicine capable of simultaneous tumor debulking and controlled drug uncaging, thereby advancing the therapeutic potential of acoustically responsive nanocarriers for solid tumor treatment.

**Figure 1.**
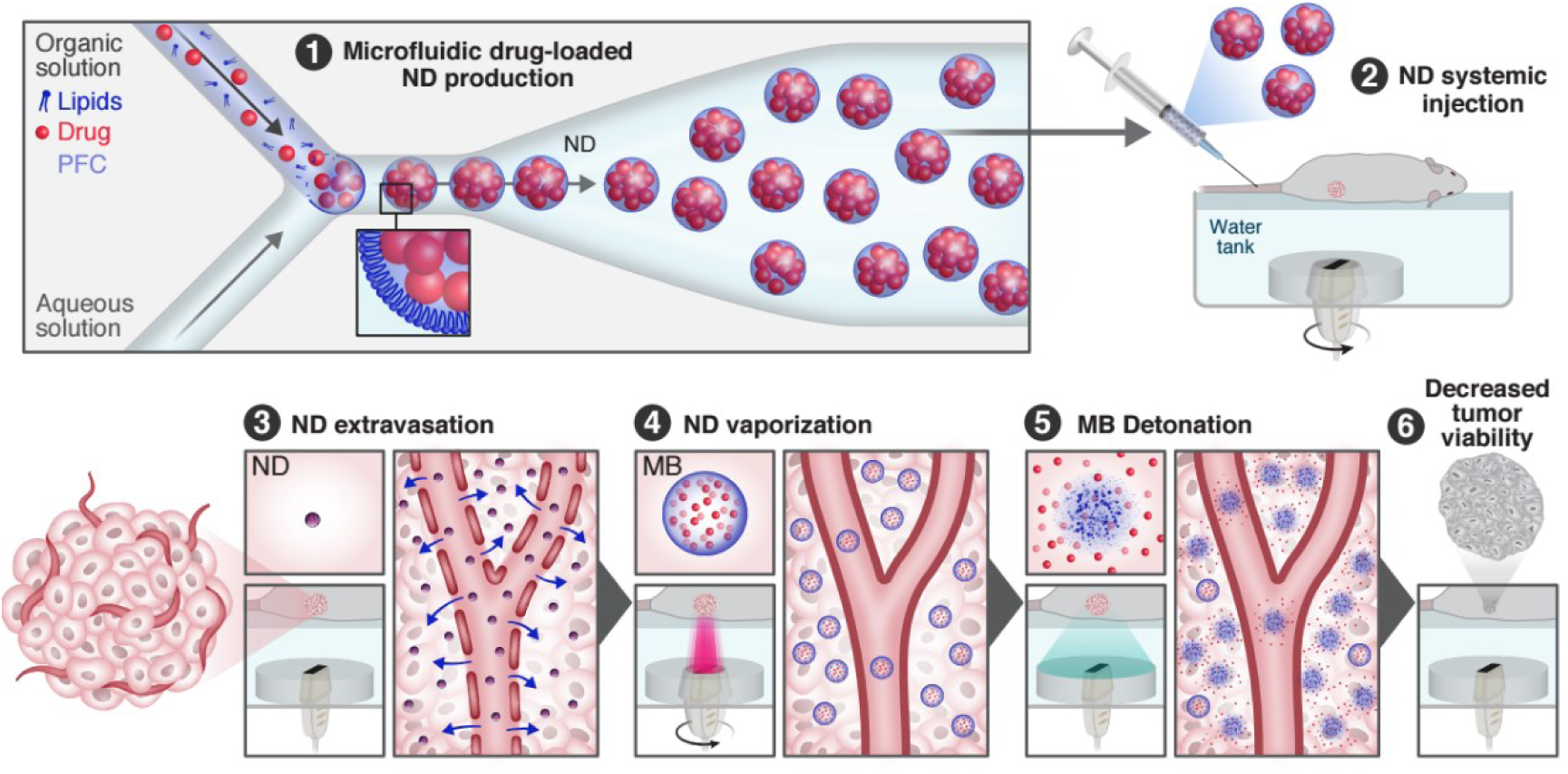
Schematic illustration of drug-encapsulated ND synthesis and dual-frequency mode US mechanotherapy. First, drug-loaded NDs are fabricated via nanoassembly microfluidic synthesis. Following systemic ND injection, a rotating imaging transducer initiates ADV to vaporize NDs into MBs. Concurrently, low-frequency therapeutic US triggers high amplitude vaporized NDs oscillations, inducing volumetric tumor tissue fractionation and drug uncaging, resulting in synergistic mechanotherapy and localized drug delivery at the tumor site.

## Results

### Ultrasound responsive nanodroplet preparation, size distribution, and stability

The first step was to optimize the microfluidic preparation method and the ND core composition. To this end, US-responsive NDs were formulated using two PFC cores, C_5_F_12_ and C_6_F_14_, via microfluidic mixing on two platforms: a glass herringbone microfluidic chip, and the NanoAssemblr system. The NanoAssemblr system operates at higher total flow rates (TFR), typically 4-24 mL/min, whereas the glass herringbone chip is generally limited to lower flow rates (0.1-5 mL/min), making the NanoAssemblr more suitable for scaled-up production and formation of smaller NDs. NDs were generated at a fixed aqueous to organic flow rate ratio (FRR) of 3:1, while varying the TFR to modulate particle size (100-2750 µL/min for the Herringbone Mixer Glass Chip and 12 mL/min for the NanoAssemblr device). This strategy enabled systematic tuning of ND size and concentration as a function of TFR. Nanoparticle Tracking Analysis (NTA) was used to quantify ND size distributions and concentrations (Table 1).

**Table 1.**
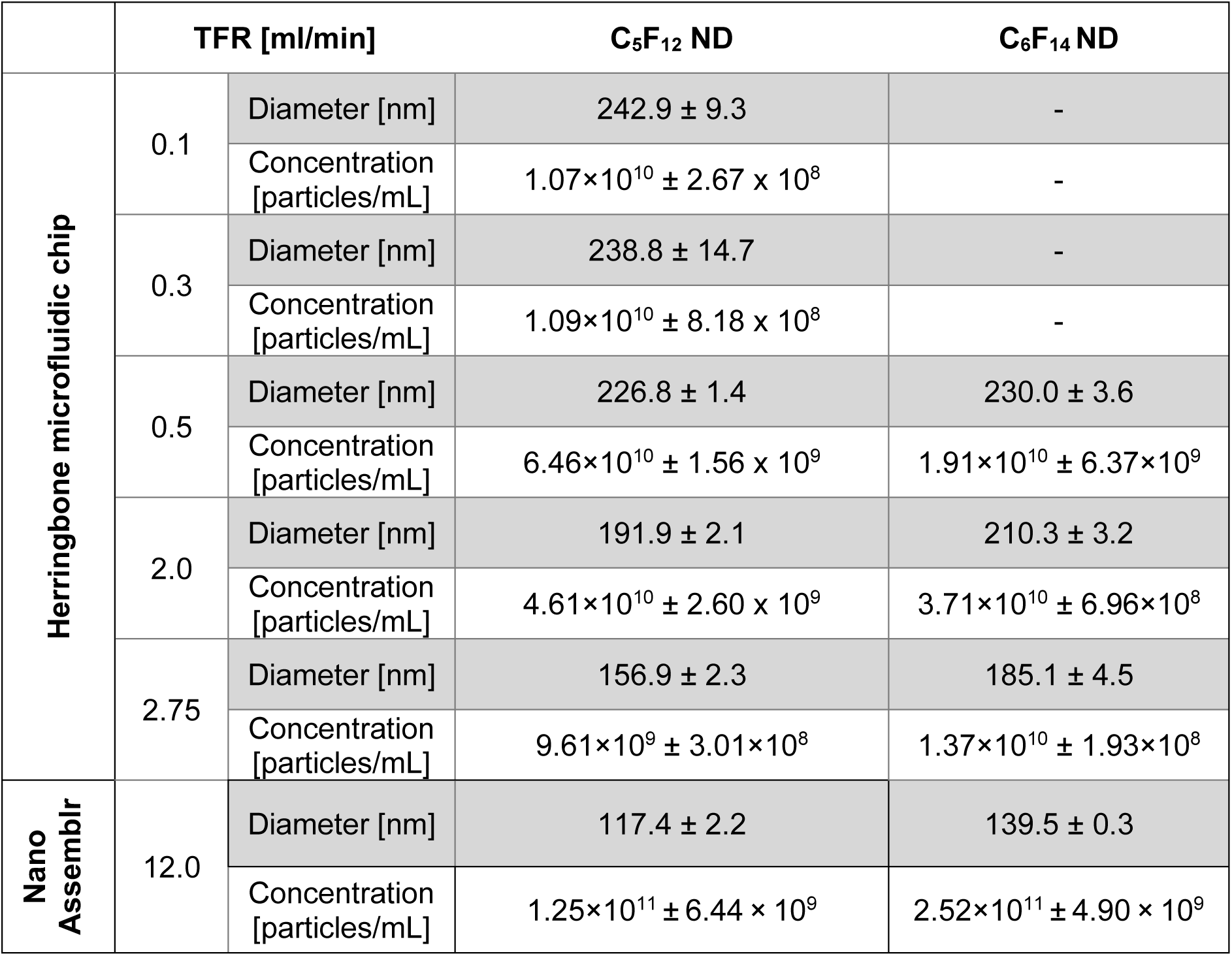
Optimization of size distribution and concentration of NDs measured by NTA.

At a high TFR of 12 mL/min on the NanoAssemblr platform with FRR = 3:1, C_5_F_12_ and C_6_F_14_ NDs exhibited mean diameters of 117.4 ± 33.4 nm and 139.5 ± 34.8 nm, respectively, with corresponding particle concentrations of 1.25 × 10^11^ ± 6.44 × 10^9^ and 2.52 × 10^11^ ± 4.90 × 10^9^ particles/mL (Figure 2c). Lower TFR (2.00-2.75 mL/min) using the Herringbone Mixer Glass Chip generated larger NDs (by ∼40-50% compared to NanoAssemblr at 12 mL/min) and lower particle concentrations across both PFC cores (Table 1). Under identical TFR conditions, C_5_F_12_ consistently produced smaller NDs than C_6_F_14_ (by ∼25-35% across TFRs tested). TEM and E-SEM imaging confirmed spherical morphology for both NanoAssemblr-generated formulations (TEM: Figure 2a,b; E-SEM: Supplementary Figure S1a,b). Zeta potential measurements revealed superior colloidal stability for C_6_F_14_ NDs (−29.8 ± 3.7 mV) compared to C_5_F_12_ NDs (−20.6 ± 3.0 mV), attributable to differences in shell-core interactions and surface charge density.Given the smaller size achievable with the NanoAssemblr, all subsequent in vitro and in vivo experiments were conducted using NanoAssemblr-fabricated NDs.

**Figure 2.**
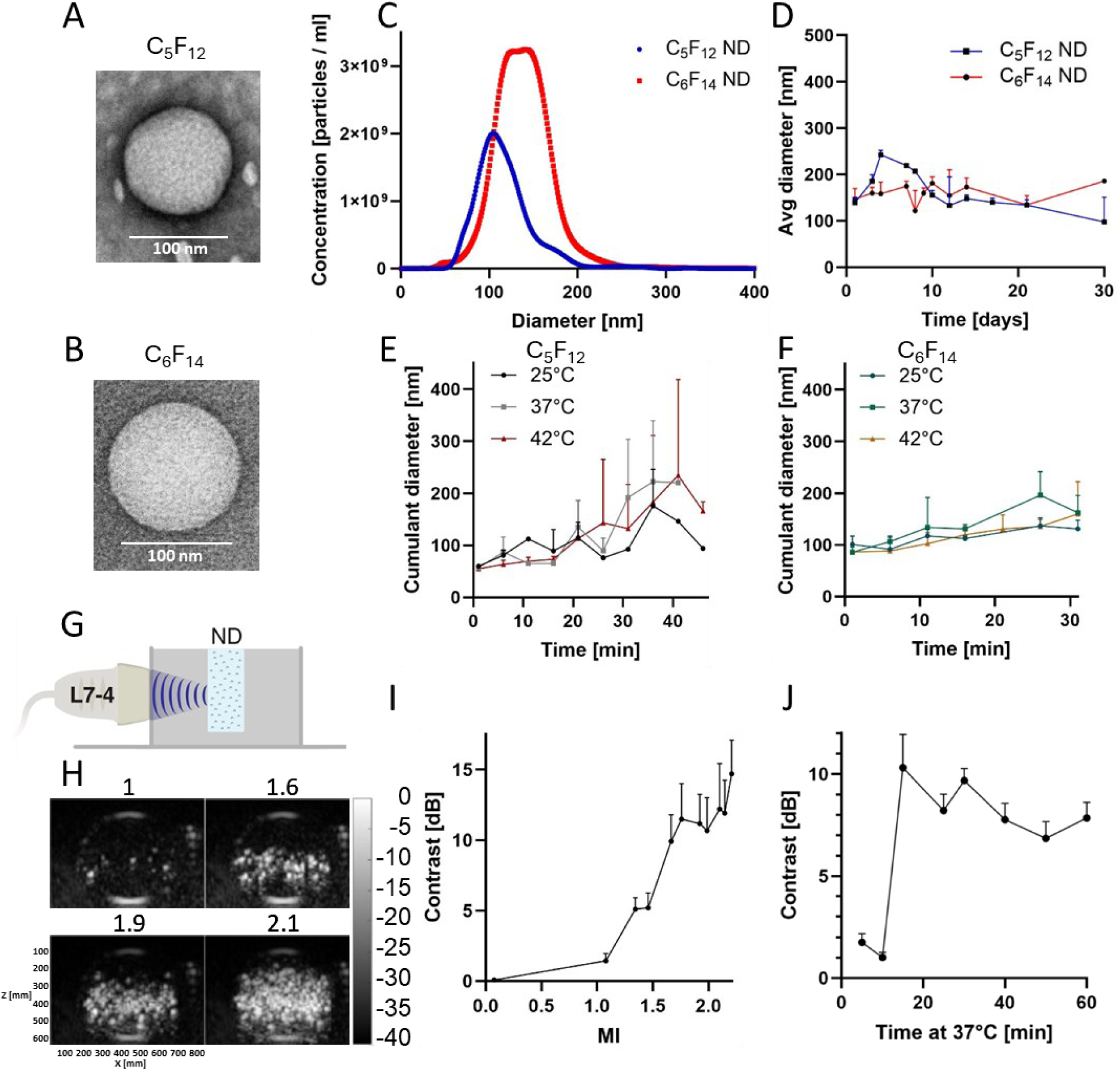
ND physicochemical and acoustic response characterization. (A, B) Transmission electron microscopy images showing the spherical morphology of (A) C_5_F_12_, (B) C_6_F_14_ NDs. Scale bars: 100 nm. (C) Nanoparticle tracking analysis (NTA) size distribution profiles comparing the hydrodynamic diameter and particle concentration of C_5_F_12_ (blue) versus C_6_F_14_ (red) NDs. (D) Long-term colloidal stability of C_5_F_12_ and C_6_F_14_ NDs stored at 4°C for 30 days. Measurements were performed at room temperature using dynamic light scattering (DLS). Data represents mean diameter ± SD (n = 3). (E, F) Thermal stability analysis via DLS, monitoring hydrodynamic size variations at ambient (25°C), physiological (37°C) and hyperthermic (42°C) temperatures for (E) C_5_F_12_, and (F) C_6_F_14_ NDs. (G) Schematic of the US activation experimental setup employing an L7-4 linear array transducer (5.5 MHz). C_6_F_14_ NDs (diluted 1:100 in degassed PBS) were injected into a cylindrical inclusion within an agarose tissue-mimicking phantom to assess acoustic droplet vaporization thresholds. (H) Representative B-mode US images illustrating pressure-dependent ADV of C_6_F_14_ NDs at 37°C. Panels show: MI 1.0 (no vaporization, baseline), MI 1.6 (onset of vaporization, sparse MBs), MI 1.9 (robust vaporization, dense MB cloud), and MI 2.1 (maximal vaporization intensity). (I) Quantitative contrast enhancement (dB) of C_6_F_14_ NDs as a function of MI ranging from 1.0 to 2.1 at 37°C. A distinct vaporization threshold is observed at MI ∼1.6. (J) Temporal stability of vaporization-induced contrast for C_6_F_14_ NDs maintained at 37°C under continuous insonation at MI 1.9, showing sustained MB persistence over time.

Next, stability tests of the two formulations were carried out over 30 days using dynamic light scattering (Figure 2d). C_6_F_14_ NDs exhibited superior stability (Figure 2d), with average size deviating by up to +26% from baseline (147.2 ± 18.1 nm at day 1). In contrast, C_5_F_12_ NDs were less stable (Figure 2d), with size increasing by up to +75% by day 5 and decreasing by −30% by day 30 relative to day 1. This can be attributed to their lower boiling point and greater susceptibility to Ostwald ripening or spontaneous phase transitions under storage conditions. Thermal stability was additionally assessed at 25°C, 37°C, and 42°C, mimicking physiological temperatures range for up to 30 minutes (Figure 2e,f). C_6_F_14_ NDs remained stable, whereas C_5_F_12_ NDs exhibited non-monotonic size fluctuations, consistent with their lower boiling point. Long-term assessments after 60 days (Supplementary Figure S1e,f) confirmed the superior stability of C_6_F_14_ NDs across all temperatures. Based on their improved storage and thermal stability, C_6_F_14_ NDs were selected for all subsequent experiments.

### Acoustic activation and vaporization behavior of NDs

NDs were next evaluated for acoustic responsiveness and vaporization thresholds. Tissue-mimicking agarose phantom experiments (Figure 2g) quantified C_5_F_12_ and C_6_F_14_ ND stability and activation behavior under thermal and acoustic conditions using an L7-4 imaging transducer (5.5 MHz). At 25°C, none of the ND formulations exhibited detectable activation or bubble formation, even at the maximum tested MI of 2.1 (Supplementary Figure S2a-d), confirming the stability of NDs at ambient temperature. For C_5_F_12_ NDs, spontaneous activation was observed at 37°C under low mechanical index (MI 0.07, Supplementary Figure S2e), and as expected, increasing the MI to 1.9 produced a corresponding increase in contrast intensity (Supplementary Figure S2f). Under physiological conditions (37°C), vaporization of C_6_F_14_ ND initiated at an MI of 1, with US echo intensity increasing monotonically with acoustic pressure (Supplementary Figure S3). This pressure-dependent enhancement yielded mean intensity gains of 5±0.8 dB at MI 1.5, 10±1.7 dB at MI 1.6, 12±3.2 dB at MI 1.9, and 14±2.4 dB at MI 2.1, with the highest contrast generation achieved at MI ≥ 1.9, corresponding to a marked 14 dB increase in US echo intensity (Figure 2h,i). These results define an activation window between MI 1.58 and 1.9 at 37°C for controlled ADV under physiological conditions.

At an elevated temperature (42°C), activation was observed at lower pressures for both formulations (MI 0.07, Supplementary Figure S2g-h). C_6_F_14_ NDs exhibited minimal partial activation at MI 0.07 and increased partial activation at MI 1.0 (Supplementary Figure S2i), while both C_5_F_12_ and C_6_F_14_ NDs showed enhanced activation at MI 1.9, with C_5_F_12_ demonstrating more pronounced vaporization (Supplementary Figure S2j,k). These results indicate that higher temperature facilitates a reduction in the vaporization threshold. In summary, acoustic droplet vaporization was temperature- and pressure-dependent, with C_6_F_14_ NDs demonstrating higher activation thresholds than C_5_F_12_ NDs.

To further investigate the influence of formulation parameters on ND stability and acoustic responsiveness, C_6_F_14_ NDs prepared at TFR of 1.0, 2.75, and 12 mL/min were compared at 37°C (Supplementary Figure S4). Flow rate influenced both ND size (as summarized in Table 1) and stability, with lower TFR producing larger, less stable NDs that exhibited partial vaporization even at low mechanical index. Specifically, NDs formulated at 1.0 mL/min showed partial activation at MI 0.07 (Supplementary Figure S4a) and more pronounced activation at MI 1.9 (Supplementary Figure S4b). NDs prepared at 2.75 mL/min similarly exhibited partial activation at low MI (0.07, Supplementary Figure S4c), though to a lesser extent than the 1.0 mL/min formulation, and reached full activation at MI 1.9 (Supplementary Figure S4d). In contrast, NDs formulated at 12 mL/min showed no detectable activation at MI 0.07 (Supplementary Figure S4e) and robust activation at high MI (1.9, Supplementary Figure S4f), confirming their superior stability. These results demonstrate that higher TFR (12 mL/min) yields smaller, more stable NDs with a well-defined activation threshold, supporting their selection for subsequent experiments.

Having established the stability and activation thresholds of NanoAssemblr C_6_F_14_ NDs, we next evaluated their behavior under repeated ultrasound exposure over time. The C_6_F_14_ NDs solution was maintained at 37°C for up to one hour, with ultrasound applied at mechanical index 1.9 at several-minute intervals to repeatedly activate the NDs and quantify the resulting contrast enhancement over time (Figure 2j), and revealed a rapid increase in contrast within the first 15 minutes (10.3±1.6 dB gain), followed by a stable plateau phase, indicating long-lasting gas-core MB formation and post-vaporization stability.

### Drug loading and ND functionalization

After confirming the acoustic activity of the particles, we next functionalized the C_6_F_14_ NDs by encapsulating fluorescent molecules and chemotherapeutic drugs. A summary of the physical properties of the PFC core (C_5_F_12_ and C_6_F_14_) and other materials (chemotherapy and fluorescent molecules) used for the formation of the NDs is provided in Supplementary Table S1. Fluorescently labeled NDs were used to quantify in vivo biodistribution and intratumoral penetration in a breast cancer model, whereas drug-loaded NDs enabled combined mechanotherapy and chemotherapy. Fluorescently labeled NDs utilized BODIPY derivatives, including BODIPY variants spanning green (BODIPY 493/503; BDPg), red (BDP TR-X-NHS ester; BDPr), and far-red (BDP 630/650-X-NHS ester; BDPfr) emission wavelengths to enable multichannel fluorescence detection and mechanistic studies. This fluorophore class provides high quantum yield and photostability^59,60^, with fluorinated side groups enabling direct C_6_F_14_ core dissolution and stable encapsulation without size increase. Chemotherapy NDs encapsulated 5-fluorouracil (5-FU). These materials were selected for their fluorinated structures, low aqueous solubility, and established utility in imaging/chemotherapeutic applications. Size, stability, zeta potential, and phase-change behavior were characterized across formulations. All encapsulated NDs exhibited comparable size distributions via NTA (Table 2), confirming successful core loading without compromising ND uniformity or microfluidic tunability. Across BODIPY, 5-FU loaded, and blank formulations, NDs showed average diameters of ∼140-160 nm, concentrations of 1.2-2.5 x 10^11^ particles/mL (Figure 3b,c), and zeta potentials between −21 and −36 mV (blank NDs ∼-30 mV), indicating robust colloidal stability. TEM confirmed well-defined core-shell nanostructures (Figure 3a), while fluorescence microscopy demonstrated minimal residual BODIPY in the supernatant after dialysis (Figure 3e), consistent with efficient core encapsulation.

**Figure 3.**
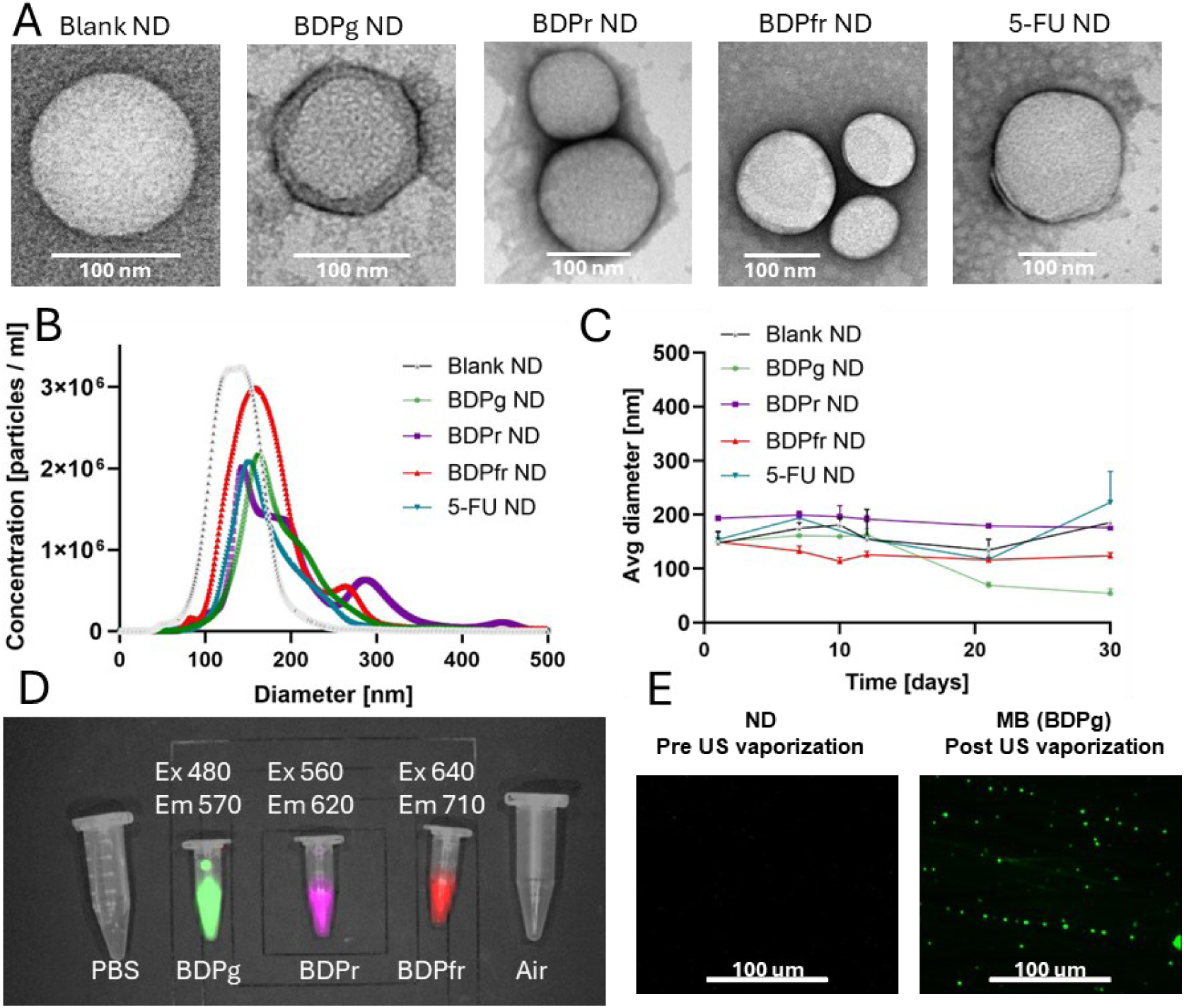
Morphology, stability, and optical properties of BODIPY-encapsulated NDs. (A) Transmission electron microscopy of blank NDs, fluorophores-loaded ND (BDPg, BDPr, BDPfr) and chemotherapy (5-FU) ND. All formulations maintain a spherical structure post-encapsulation. Scale bar: 100 nm. (B) Comparative NTA profiles displaying the size distribution and particle concentration for blank, dye-loaded, and drug-loaded NDs. (C) Long-term stability analysis tracking the average hydrodynamic diameter of blank and encapsulated NDs over a 30-day storage period. (D) IVIS fluorescence imaging of Eppendorf tubes containing PBS (control), air (control), and the three BODIPY-loaded ND variants. The panel highlights successful spectral separation and distinct fluorescence emission for BDPg (Ex 480 nm/Em 570 nm), BDPr (Ex 560 nm/Em 620 nm), and BDPfr (Ex 640 nm/Em 710 nm) NDs. (E) Fluorescence microscopy of BDPg NDs before and after US exposure. The Pre-US state shows no detectable fluorescence structures, while the Post-US state reveals the formation of green-fluorescent MBs. Scale bar: 100 µm.

**Table 2.** Size distribution, concentration and zeta potential of blank, BODIPY and 5-FU loaded NDs.

| NDs type | Average size [nm] | Concentration [particles/mL] × 10<sup>11</sup> | Zeta potential [mV] |
| --- | --- | --- | --- |
| Blank ND | 139.5 ± 0.3 | 2.52 ± 0.05 | -29.80 ± 3.70 |
| BDPg ND | 162.3 ± 58.6 | 1.34 ± 0.11 | -21.19 ± 0.96 |
| BDPr ND | 143.3 ± 79.0 | 1.53 ± 0.02 | -21.48 ± 4.54 |
| BDPfr ND | 158.5 ± 60.7 | 2.11 ± 0.03 | -36.33 ± 1.31 |
| 5-FU ND | 151.0 ± 45.5 | 1.18 ± 0.03 | -23.07 ± 0.33 |

BODIPY and 5-FU loaded NDs exhibited distinct 30-day stability profiles (Figure 3c) under refrigerated storage (4°C; measurements at 25°C). BDPfr NDs (initial 193.5 ± 1.4 nm) demonstrated long-term stability, maintaining diameters within ∼8% of initial size through day 30. 5-FU NDs (initial 154.0 ± 13.7 nm) showed moderate stability with fluctuations up to ±45% relative to baseline over 30 days. In contrast, BDPg NDs (initial 149.4 ± 3.4 nm) were less stable, exhibiting a progressive size reduction of ∼65% by day 30.

Multispectral IVIS analysis characterized the excitation and emission properties for each BODIPY-loaded ND formulation. BDPg NDs exhibited peak fluorescence at 480/570 nm, BDPr NDs at 560/620 nm, and BDPfr NDs at 640/710 nm (Figure 3d). These optimized spectral pairs provided maximal signal intensity for each type while maintaining separation between formulations. Specificity was further validated by the absence of signal in control samples (PBS and air blanks).

### ADV and cavitation triggered drug release

Next, we sought to evaluate the ability of the NDs to achieve controlled release of the fluorescent payload. First, we assessed the solubility of free BODIPY (BDPg) in different solvents (Figure 4a), demonstrating that BODIPY remained undissolved in PBS solution but readily dissolved in C_6_F_14_ PFC solution, forming a homogeneous green-yellow solution. Based on this solubility profile, we formulated BODIPY-loaded NDs by dissolving the fluorophore in C_6_F_14_ and encapsulating the solution within a lipid shell. Then, we verified that the ADV capability was preserved. Upon US activation (L7-4 transducer, 5.5 MHz, MI 1.9) of BODIPY NDs ADV of the C_6_F_14_ core generated fluorescent MBs, visualized by microscopy (Figure 3e, pre/post-vaporization). Pre-activation NDs (<200 nm) were undetectable at 20x magnification (100 µm scale bar), whereas post-ADV MBs exhibited pronounced fluorescence enhancement attributable to ADV-induced volume expansion. Similar results were obtained with red BODIPY NDs with fluorescent MBs formation post-activation (Supplementary Figure S1c,d). To induce payload release, a two-step activation-detonation process was applied using an US guided focused US system. First, ND vaporization was induced using a rotating imaging array operating at a center frequency of 3.5 MHz and MI = 1.67 to enable volumetric ND activation. Subsequently, collapse of the vaporized NDs was driven using a therapeutic transducer operating at a center frequency of 105 kHz and MI = 0.84. Applying this sequence to BDPg NDs triggered substantial payload release (Figure 4b), quantified by UV-Vis spectrophotometry. Due to the high density of C_6_F_14_(1.68*g/ml*) relative to the aqueous medium, NDs sedimented, enabling supernatant analysis of free fluorophore. Calibration against BDPg standards (0.1-0.6 µM, Figure 4c) revealed a 2.8-fold fluorescence increase post-treatment (0.61 ± 0.11 µM supernatant vs. 0.22 ± 0.08 µM pre-treatment, Figure 4d), confirming efficient ADV and cavitation-triggered release with this dual-frequency paradigm. After confirming the ability of the NDs to release fluorescent molecules, we next evaluated their capacity to encapsulate chemotherapeutic drugs. First, we evaluated the solubility of 5-FU in C_6_F_14_ compared to DI water using UV-Vis spectrophotometry. Full-spectrum scanning (λ_ex_ 240-295 nm, λ_em_ 310 ± 5 nm) revealed a distinct absorbance profile for 5-FU in C_6_F_14_, characterized by a sharp intensity peak around 270 nm, significantly exceeding the signal observed in DI water (Figure 4e). Quantitative analysis was subsequently performed using the optimized parameters (λ_ex_ 265 ± 20 nm, λ_em_ 310 ± 20 nm), where 5-FU dissolved in C_6_F_14_ exhibited a ∼6-fold increase in fluorescence intensity compared to 5-FU in DI water (Figure 4f). Control measurements of neat solvents showed baseline intensities of 15,910 ± 384 a.u. for C_6_F_14_ and 7,510 ± 79 a.u. for DI water. When corrected for solvent background, the net signal for 5-FU in C_6_F_14_ (34,758 a.u.) was 31-fold higher than in DI water (1,106 a.u.), confirming the superior solubility and retention of fluorinated 5-FU molecules within the PFC phase.

**Figure 4.**
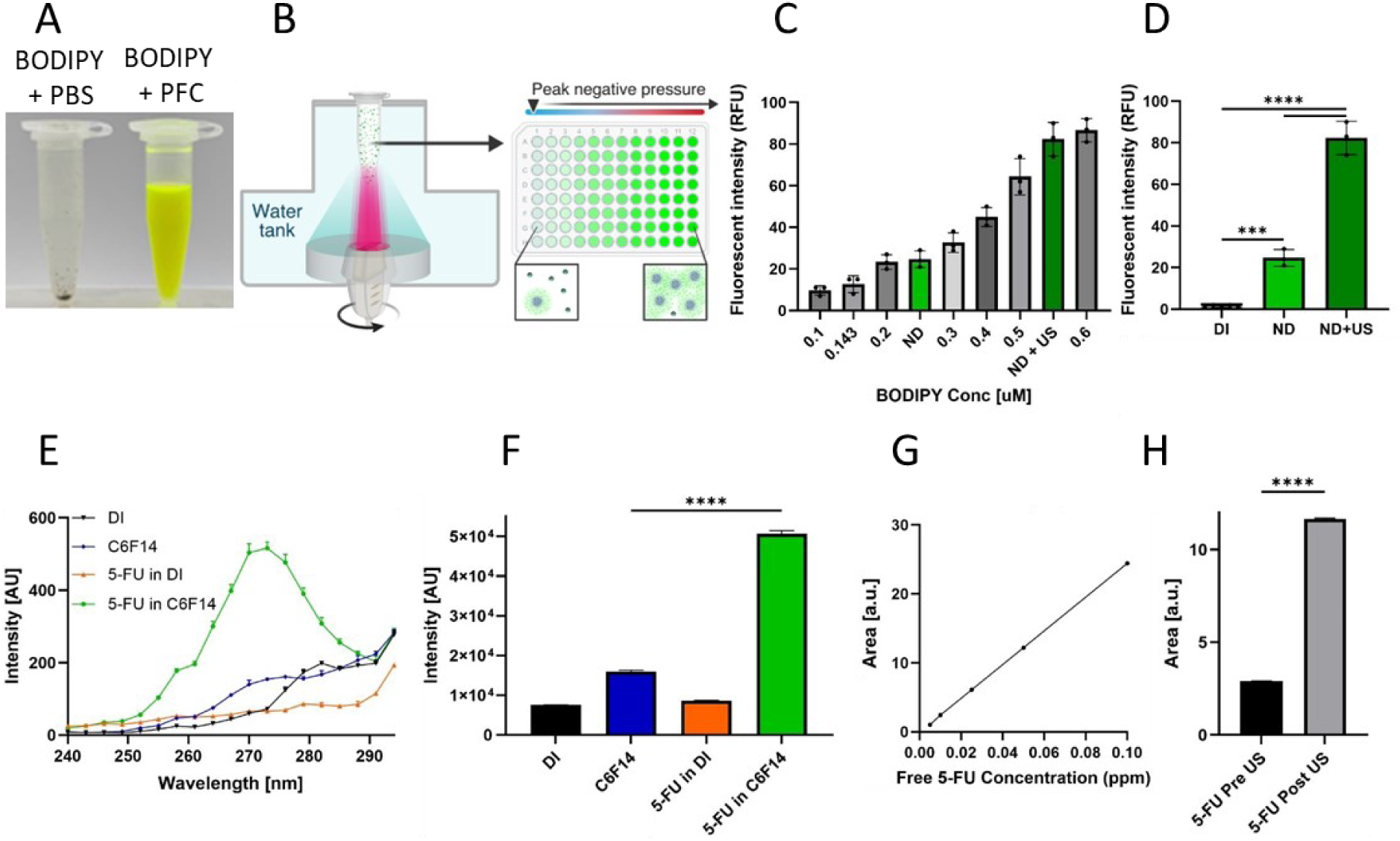
Ultrasound-triggered payload release kinetics and validation of core-loading solubility. (A) Solubility comparison of BODIPY in different solvents. Left: BODIPY in PBS solution remains undissolved, forming a solid precipitate at the bottom of the tube. Right: BODIPY in PFC (C_6_F_14_) demonstrates complete dissolution, yielding a homogeneous green-yellow solution. (B) Schematic illustration of the in vitro release experimental setup. Payload release was triggered using a simultaneous dual-frequency US system (3.5 MHz imaging transducer, MI 0.84, 105 kHz therapy transducer, MI 1.67). Release efficiency was quantified by analyzing the fluorescence/absorbance of the supernatant fraction collected before and after US exposure. (C) Quantification of BDPg release showing the calibration curve (fluorescence intensity vs. concentration, R^2^=0.979) overlaid with supernatant measurements. A ∼4-fold increase in fluorescence intensity is observed in the post-US supernatant compared to pre-US controls. (D) Comparative fluorescence analysis of top supernatant fractions from DI water (background), intact NDs (passive leakage), and US-activated NDs (active release). (E) Spectroscopic characterization of 5-FU solubility. Representative excitation spectra (Em = 310 nm) of 5-FU dissolved in C_6_F_14_ versus DI water, revealing a characteristic intensity peak at ∼270 nm specific to the fluorinated phase. (F) Quantification of 5-FU fluorescence intensity (Ex = 265 ± 20 nm, Em = 310 ± 20 nm) in C_6_F_14_ and DI water compared to pure solvent controls. The significant signal enhancement confirms the successful dissolution and core-loading of 5-FU within the C_6_F_14_ matrix. (G) HPLC calibration curve correlating peak area with free 5-FU concentration, used to calculate the release concentrations in panel H (R^2^=0.99). (H) HPLC quantification of 5-FU release efficiency, showing a significant increase in the integrated peak area (a.u.) for post-US samples compared to pre-US baselines (****p < 0.0001). Data represent mean ± SD (n = 3). Statistical significance was calculated using one-way ANOVA with Tukey’s multiple comparison test (*p < 0.05, **p < 0.01, ***p < 0.001, ****p < 0.0001).

US-triggered 5-FU release experiments were then performed as described for the BODIPY-loaded NDs. The released drug was quantified by HPLC using a calibration curve generated from free 5-FU (0.005-0.1 ppm, R^2^ = 0.9861, Figure 5g). For analysis, 10 µL of supernatant was collected from the top of each 1 mL sample before and after US application, diluted 100-fold, filtered, and processed for HPLC. These measurements confirmed an increase in 5-FU concentration post-sonication. At a flow rate of 0.8 *mL/min* with a 50 µL injection volume, the chromatographic peak area increased from 2.91 to 11.65 (Figure 5h), representing a 4.0-fold increase. This translated to original solution concentrations rising from 1.2 ppm (pre-US) to 4.8 ppm (post-US), after correction for dilution.

**Figure 5.**
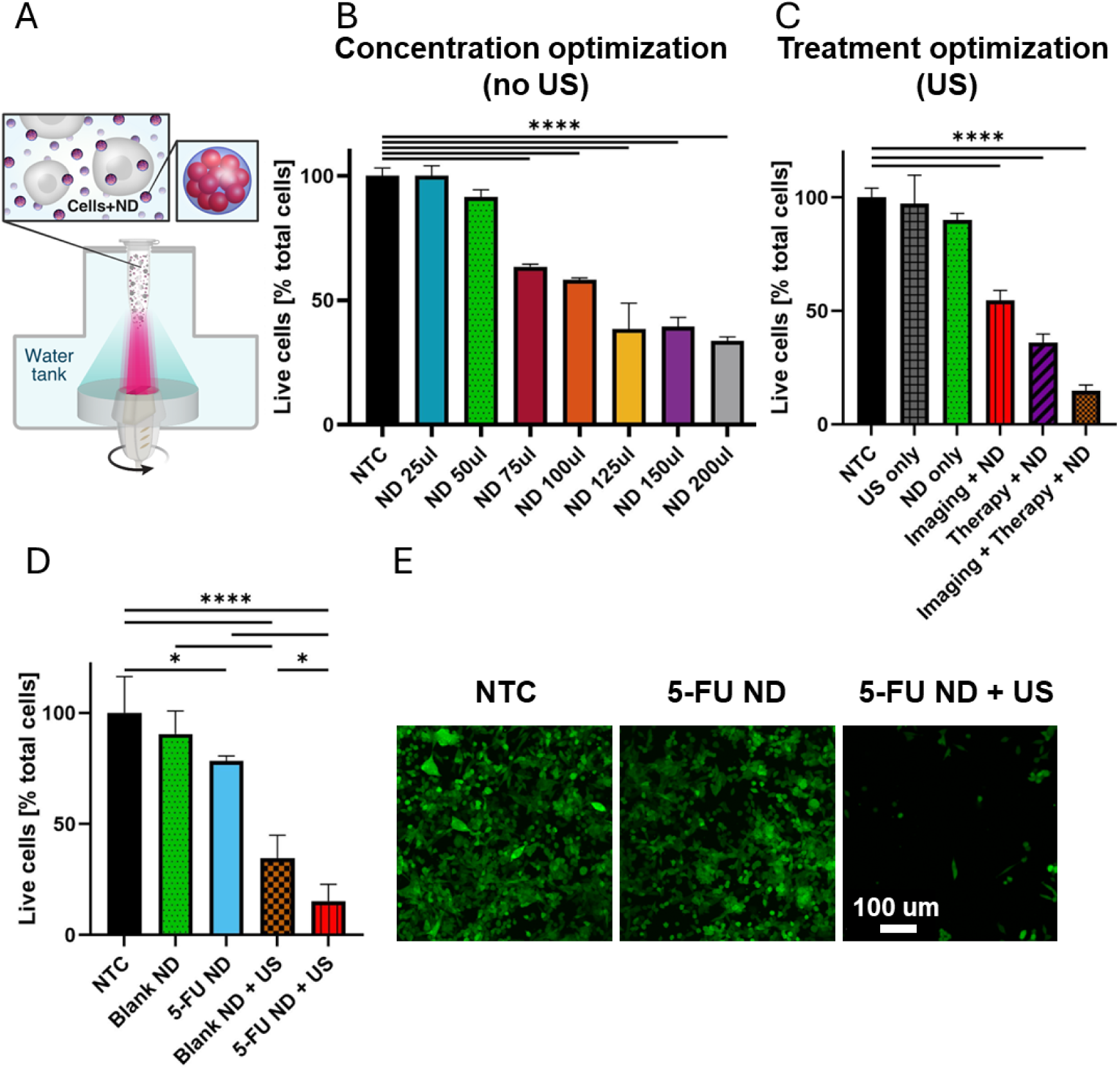
In vitro optimization and therapeutic efficacy of ND-mediated therapy.(A) Schematic representation of the in vitro experimental setup. An Eppendorf containing a suspension of cancer cells mixed with NDs was positioned at the acoustic focal point of the US-guided focused US system and exposed to US. Cell viability was assessed 72 h after treatment. (B) Effect of ND concentration on cell viability without ultrasound exposure. (C) Comparative analysis of cell viability of NTC, dual-frequency US only, NDs only, imaging transducer + NDs, therapy transducer + NDs, and the combined dual-frequency + NDs. (D) Quantitative viability analysis of live cells across therapeutic groups: NTC, blank ND, 5-FU ND, blank ND + US, and 5-FU ND + US. The combined treatment demonstrates significant synergistic cytotoxicity. (E) Representative confocal microscopy images of GFP-expressing 4T1 breast cancer cells following selected treatments. Scale bar: 100 µm. Data represents mean ± SD (n = 6). Statistical significance was calculated using one-way ANOVA with Tukey’s multiple comparison test (*p < 0.05, **p < 0.01, ***p < 0.001, ****p < 0.0001).

### In vitro evaluation of ND-mediated mechanotherapy and drug uncaging

In vitro experiments were conducted in breast cancer cells. First, we optimized the effect of blank NDs alone on cell viability. Cells were incubated with NDs for 72 h, and cell viability was measured. The NDs concentration was 2.52 × 10⁸ NDs/µL, and the ND volume varied between 25-200 µL. A dose-dependent cytotoxicity was observed (Figure 5b). Low concentrations (25-50 µL) maintained >90% viability comparable to untreated controls (NTC; *p > 0.05), whereas a ND volume larger than 125 µL resulted in >60% cell death. These results are consistent with prior studies^43,61^. Based on these results, 50 µL of blank NDs was selected as the optimal concentration for subsequent in vitro US experiments.

Before evaluating drug delivery effects, US-triggered mechanotherapy was assessed using the optimized parameters and the dual-frequency setup described in the uncaging experiments. Because the activation protocol involves two sequential US steps, we first evaluated the contribution of each step to cell viability. Cell viability was therefore assessed following ND vaporization induced by the imaging transducer alone, low-frequency therapeutic US applied to NDs without prior vaporization, and the full dual-frequency protocol combining both steps. Additional control groups included US-only conditions without NDs and untreated cells. While all control groups maintained high viability (NTC: 100 ± 4.18%; US only: 97.23 ± 12.59%; blank NDs only: 90.11 ± 2.91%), ND+US combinations produced progressive reductions in viability. Imaging-frequency US with NDs reduced viability to 54.65 ± 4.38% (****p < 0.0001 vs NTC), therapeutic low-frequency US with NDs to 36.02 ± 3.86% (****p < 0.0001), and the full dual-frequency protocol to 14.82 ± 2.55% (****p < 0.0001), demonstrating a strong mechanotherapeutic effect. Prior to evaluating US-triggered 5-FU uncaging, the cytotoxic effect of free 5-FU was first assessed. Free 5-FU exhibited dose-dependent cell cytotoxicity across concentrations of 9.6-76.9 µM with a cell viability decreasing from 100% in the NTC to below 20% at 9.6 and 19.2 µM and under 10% at 38.4 and 76.9 µM (Supplementary Figure S5), establishing the baseline sensitivity of the cells to the chemotherapeutic agent. We next evaluated US-triggered 5-FU release from drug-loaded NDs and compared the resulting combined chemo-mechanical effect with the purely mechanical effect produced by blank NDs under the same ultrasound conditions (Figure 5d). The strongest cytotoxic effect was observed when US was applied to 5-FU-loaded NDs, reducing cell viability to 15.26 ± 7.61% (****p < 0.0001 vs NTC; *p < 0.05 vs blank NDs + US). In comparison, activation of blank NDs by US reduced viability to 34.49 ± 10.42%, indicating that the additional cytotoxicity arises from ultrasound-triggered drug release. Control conditions, including untreated cells, blank NDs alone, and passive 5-FU ND delivery, had only minimal impact on cell viability (Figure 5d). Confocal microscopy of GFP-expressing breast cancer cells (Figure 5e) visually confirmed the progressive cytotoxic response across the treatment groups.

### In vivo pharmacokinetics and tumor accumulation of ND

Following the demonstration of combined mechanochemical efficacy in vitro, NDs were evaluated in a mice breast cancer model to characterize their in vivo pharmacokinetics, blood circulation, and tumor accumulation prior to therapeutic studies. Systemic administration in breast cancer-bearing mice enabled quantification of ND blood clearance, biodistribution, and extravasation into solid tumors, providing a basis for selecting dosing and timing parameters for subsequent in vivo US therapeutic experiments. First, systemic pharmacokinetics and biodistribution of NDs was characterized. Although BDPg NDs exhibited peak fluorescence at 480/570 nm (Figure 3d), in vivo biodistribution results showed significant autofluorescence from mouse tissues at these wavelengths. To overcome this, BDPfr NDs, with peak excitation/emission at 640/710 nm, were selected for in vivo imaging to minimize background signal.

Blood pharmacokinetics of BDPfr NDs were quantified by IVIS fluorescence analysis of serial blood samples (10 min, 3h, 24h post-injection; Figure 6a). BDPfr NDs exhibited rapid signal increase (10 min: 2.56×10^8^ ± 9.41×10^7^ radiant efficiency vs. baseline 1.77×10^7^ ± 2.40×10^6^) followed by prolonged circulation (3h: 1.86×10^8^ ± 1.89×10^7^; 24h: 1.91×10^7^ ± 1.02×10^7^), demonstrating extended blood retention.

**Figure 6.**
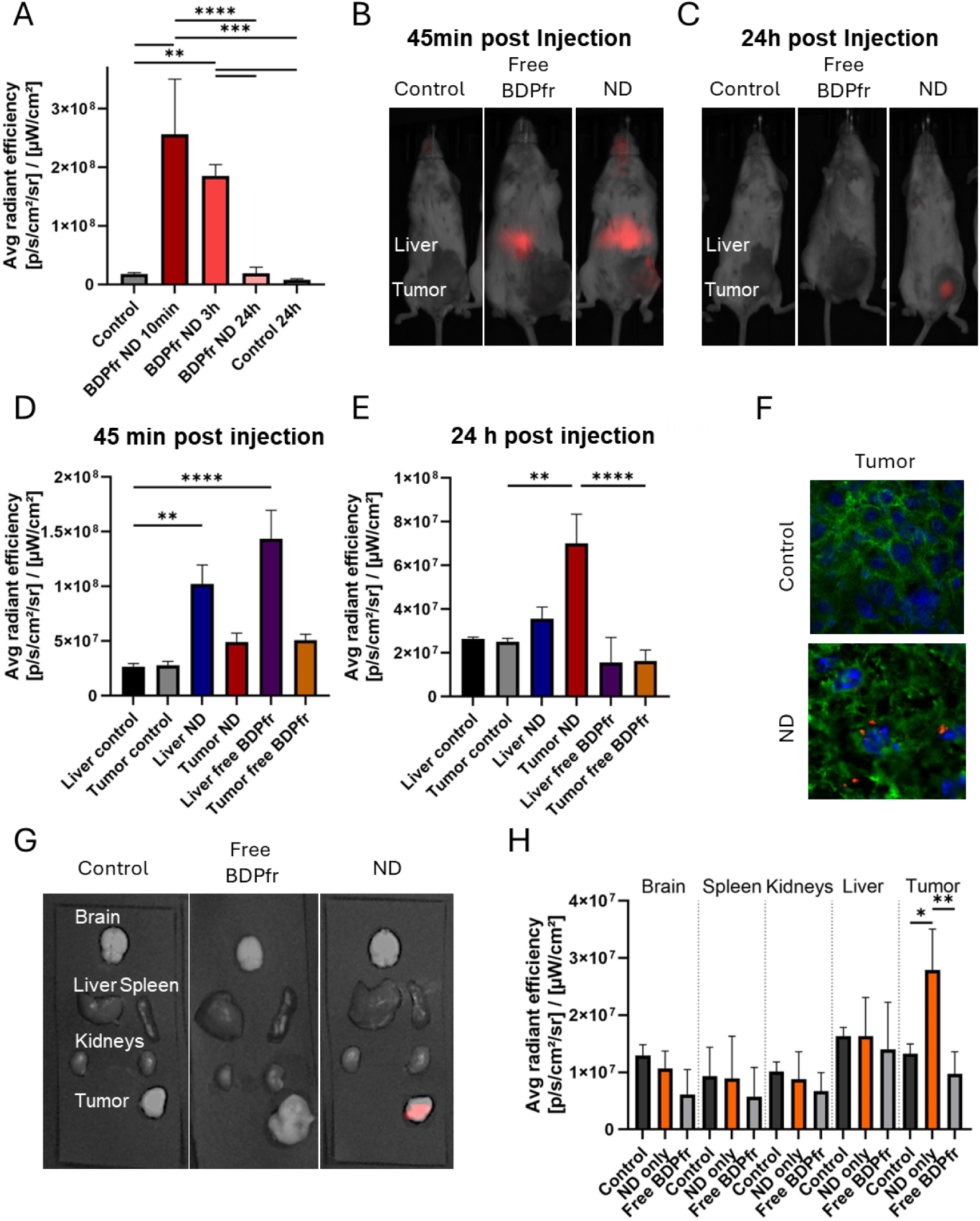
In vivo biodistribution and tumor accumulation of BDPfr NDs. (A) Time-dependent fluorescence intensity of BDPfr in blood samples 10 min, 3 h and 24 h post injection following administration of BDPfr NDs, compared with non-treated controls (NTC). (B, C) Representative whole-body IVIS fluorescence imaging of tumor-bearing mice at (B) 45 minutes, and (C) 24 hours post-injection. Groups include non-injected control, free BDPfr dye, and BDPfr NDs. Regions of interest (ROIs) indicate liver and tumor locations. (D, E) Quantitative analysis of fluorescence intensity in liver and tumor regions at (D) 45 minutes, and (E) 24 hours, highlighting statistically significant tumor retention in the ND-treated group relative to controls and free dye. (F) Confocal microscopy of histological tumor sections from control and ND-treated mice. Images display nuclei (DAPI, blue), cell membranes (WGA, green), and NDs (BDPfr ND, red). The presence of intracellular red dots confirms deep tissue penetration and cellular uptake of the NDs. (G) Ex vivo fluorescence imaging of excised organs (brain, liver, spleen, kidneys, tumor) 24 hours post-injection, validating the biodistribution patterns observed in vivo. (H) Quantification of ex vivo radiant efficiency across harvested organs 24 hours post-injection. The analysis confirms preferential accumulation and high-intensity signal retention within the tumor for the ND group compared to the control and free dye groups, demonstrating effective passive targeting via the EPR effect. Data represent mean ± SD, N =3. Statistical significance was calculated using one-way ANOVA with Tukey’s multiple comparison test (*p < 0.05, **p < 0.01, ***p < 0.001, ****p < 0.0001).

Subsequently, IVIS imaging was used to track BDPfr ND and free BDPfr dye biodistribution in breast cancer-bearing mice across 24 hours, with representative time points shown at 45 minutes and 24 hours. At 45 minutes post-injection (Figure 6b,d), ND-treated mice exhibited significantly elevated fluorescence in the liver (1.02 × 10^8^ ± 1.73 × 10^7^ average radiant efficiency) and tumor (4.91 × 10^7^ ± 8.06 × 10^6^). This represented a 3.8-fold increase in liver fluorescence compared to controls (2.65 × 10^7^ ± 2.97 × 10^6^) and a 1.8-fold increase in tumor fluorescence relative to controls (2.79 × 10^7^ ± 3.61 × 10^6^). ND-mediated accumulation was comparable to that of free dye-injected animals, which showed liver fluorescence of 1.44 × 10^8^ ± 2.60 × 10^7^ (5.4-fold above control) and tumor fluorescence of 5.08 × 10^7^ ± 5.51 × 10^6^ (1.8-fold above control). One-way ANOVA with Tukey’s multiple comparison test confirmed significant differences among group means at this time point (**p < 0.01, ****p < 0.0001). By 24 hours post-injection (Figure 6c,e), a clear divergence between formulations emerged. ND-treated mice retained elevated tumor fluorescence (7.01 × 10^7^ ± 1.34 × 10^7^), 2.8-fold above controls (2.51 × 10^7^ ± 1.56 × 10^6^) and 4.3-fold greater than free dye (1.63 × 10^7^ ± 5.03 × 10^6^). In contrast, liver fluorescence in the ND group (3.56 × 10^7^ ± 5.33 × 10^6^) and free dye group (1.56 × 10^7^ ± 1.14 × 10^7^) were comparable to control (2.64 × 10^7^ ± 8.49 × 10^5^), indicating efficient hepatic clearance. One-way ANOVA with Tukey’s multiple comparison test again indicated significant differences among group means (**p < 0.01, ****p < 0.0001).

Extended biodistribution kinetics over 90 hours post-injection (Supplementary Figure S6) further elucidated the accumulation and retention behavior of BDPfr NDs. Within minutes of administration, ND-treated mice displayed a rapid fluorescence increase in the tumor region, which remained significantly elevated and stable throughout the initial 4.5-hour imaging period (Supplementary Figure S6a). Long-term tracking revealed that while liver fluorescence peaked early and declined to near-baseline levels by 24 hours, tumor fluorescence continued to rise, reaching maximum intensity at approximately 55 hours before declining slightly by 90 hours while remaining above control levels (Supplementary Figure S6b). The kinetic profile over 220 minutes (Supplementary Figure S6c) further illustrates the distinct accumulation and retention dynamics for each experimental group, with NDs demonstrating sustained tumor localization compared to free dye. These results confirm strong and sustained tumor retention of BDPfr NDs compared to transient hepatic uptake, supporting their potential for targeted drug delivery applications.

To validate these in vivo observations, ex vivo analysis of harvested organs was performed at 24 hours post-injection (Figure 6g,h). This confirmed maximal tumor accumulation for NDs (2.79 × 10^7^ ± 7.14 × 10^6^), which was significantly higher than free dye (9.71 × 10^6^ ± 3.88 × 10^6^) and control (1.32 × 10^7^ ± 1.71 × 10^6^). This represented a 2.9-fold increase compared to free dye and a 2.1-fold increase relative to control. In other organs, ND-associated fluorescence was generally similar to background, except in spleen and kidneys, where values and variances were comparable to controls (Brown-Forsythe tests, all p > 0.22), indicating no significant off-target accumulation. One-way ANOVA with Tukey’s multiple comparison test for ex vivo data validated significant differences among tumor ND group with tumor control (*p < 0.05) or with tumor free dye group (**p < 0.01).

To determine whether this macroscopic tumor retention of BDPfr NDs reflects deep tissue penetration and cellular uptake, we next examined tumor sections using confocal microscopy. Tumors collected 24 h post-injection were processed for high-resolution fluorescence imaging. Confocal microscopy (Figure 6f) showed penetration of BDPfr NDs within tumor sections, with red fluorescence (BDPfr ND) localized adjacent to cell nuclei (blue, DAPI) across multiple regions. Co-staining with WGA Alexa Fluor 488 (green) highlighted cell membrane architecture and confirmed intracellular localization of NDs in merged images.

### In vivo mechanotherapy in a breast cancer mouse model

To isolate mechanotherapy effects prior to combined treatment evaluation, blank NDs were tested with dual-frequency US (Figure 7a). 10 minutes post-systemic blank ND administration (110 μL per mouse), mice received 2 minutes of dual-frequency US. Histological analysis of tumor sections harvested 24 hours post-treatment revealed cavitation-induced lesions in the NDs + dual-frequency group. In contrast, control groups, including untreated mice, US alone, or NDs alone, exhibited minimal to no histological evidence of damage (mean lesion areas: 1.55 ± 0.92% for untreated, 3.42 ± 1.39% for US only). While the dual-frequency ND-mediated group demonstrated significantly greater tissue fractionation, with lesion areas averaging 16.32 ± 5.02% (Figure 7b,d, ***p < 0.001). Following ND-induced mechanotherapy, we hypothesized that the resulting vascular disruption would increase tumor vascular permeability and thereby enhance subsequent ND accumulation. To test this, BDPfr NDs were systemically injected 24 h after FUS treatment, and organ accumulation was quantified ex vivo using IVIS imaging (Figure 7c). Tumors in the BDPfr ND-only group (2.79×10^7^ ± 7.14×10^6^ radiant efficiency) showed an 2-fold increase relative to NTC (1.32×10^7^ ± 1.71×10^6^). Prior mechanotherapy (Blank ND + US) further increased tumor fluorescence to ∼5-fold above control (6.44×10^7^ ± 1.27×10^7^). The highest signal was observed in tumors pre-treated with combined mechanochemotherapy (5-FU ND + US, 8.48×10^7^ ± 1.54×10^7^), corresponding to ∼7-fold enrichment relative to NTC and ∼3-fold relative to BDPfr ND alone. These results indicate that US-mediated mechanical disruption, particularly when combined with chemotherapeutic delivery, increases tumor vascular permeability and promotes intratumoral ND accumulation.

**Figure 7.**
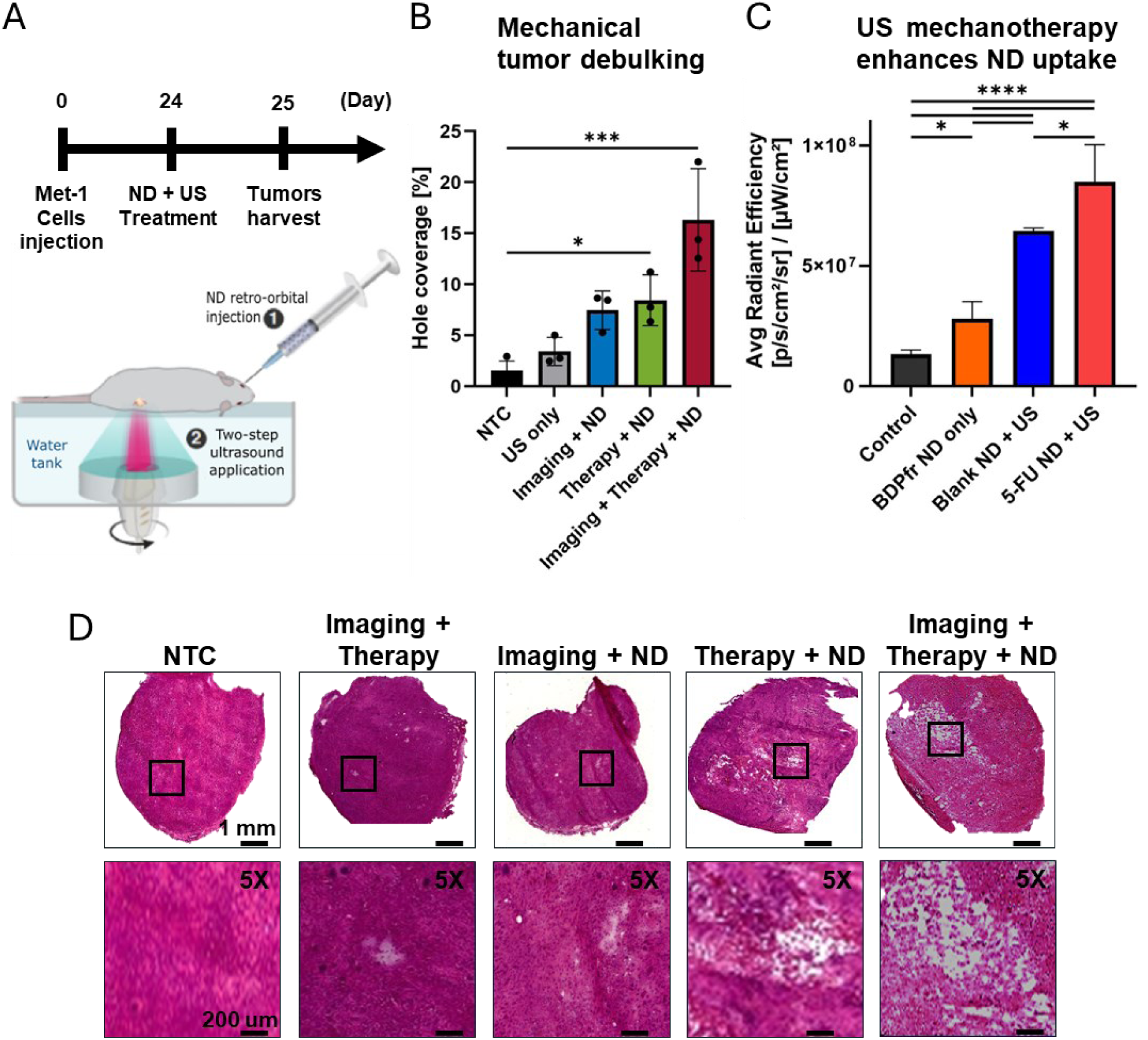
Two-step US applied to blank NDs includes mechanotherapy in vivo in a breast cancer tumor model. (A) Illustration of the in vivo treatment protocol and timeline. NDs were administered systemically, followed by dual-frequency insonation of the tumor to induce mechanotherapy. (B) Quantification of histological damage, expressed as percentage of hole coverage (fractionated tissue area) across treatment groups: NTC, US only, imaging US + ND, therapy US + ND, and the full protocol (imaging + therapy + ND). The combined regimen induced maximal tissue fractionation. (C) Ex vivo IVIS analysis of average radiant efficiency in tumors following subsequent injection of BDPfr NDs. Groups include: naïve tumor control, tumor with BDPfr ND only, tumor pre-treated with blank ND + US, and tumor pre-treated with 5-FU ND + US. Enhanced fluorescence was observed in tumors pre-treated with mechanical ablation (blank ND + US), with significantly greater accumulation following mechanotherapy (5-FU ND + US), indicating that therapeutic disruption progressively enhances vascular permeability and drug delivery. (D) Representative histological photomicrographs (H&E staining) of the treated tumors. Groups include NTC, US only, imaging + ND, therapy + ND, and the combined dual-frequency + ND activation. Black squares indicate regions of successful tumor fractionation (mechanical ablation). Scale bars: 1 mm (whole tumor cross-sections), 200 µm (5× magnification). Data represent mean ± SD, N=3. Statistical significance was calculated using one-way ANOVA with Tukey’s multiple comparison test (*p < 0.05, **p < 0.01, ***p < 0.001, ****p < 0.0001).

### Combined mechanotherapy and chemotherapy drug uncaging in vivo

Lastly, the combined mechanotherapy and chemotherapy uncaging therapeutic effect was assessed by monitoring tumor growth and survival compared to four groups receiving weekly interventions: NTC, 5-FU ND alone, blank ND + US, and 5-FU ND + US. Tumor growth was monitored over four weeks to evaluate the therapeutic effect of combined mechanotherapy and chemotherapy uncaging (Figure 8a,b). Baseline tumor volumes (Figure 8b) were comparable across groups (NTC: 117.6 ± 71.75 mm^3^; 5-FU ND: 81.93 ± 32.38 mm^3^; blank ND + US: 122.3 ± 51.30 mm^3^; 5-FU ND + US: 92.98 ± 43.67 mm^3^). Over the treatment period, NTC and 5-FU ND alone exhibited rapid tumor progression, reaching mean volumes of 768.2 ± 581.5 mm^3^ and 934.0 ± 417.8 mm^3^, respectively. In contrast, blank ND + US produced moderate tumor growth inhibition (528.1 ± 376.6 mm^3^), while the combined treatment (5-FU ND + US) showed the strongest effect, limiting tumor growth to 265.6 ± 99.48 mm^3^ at four weeks (Figure 8a). This represented a 2.0-fold smaller tumor volume compared to blank ND + US, a 2.9-fold reduction relative to NTC controls, and a 3.5-fold decrease compared to 5-FU ND alone.

**Figure 8.**
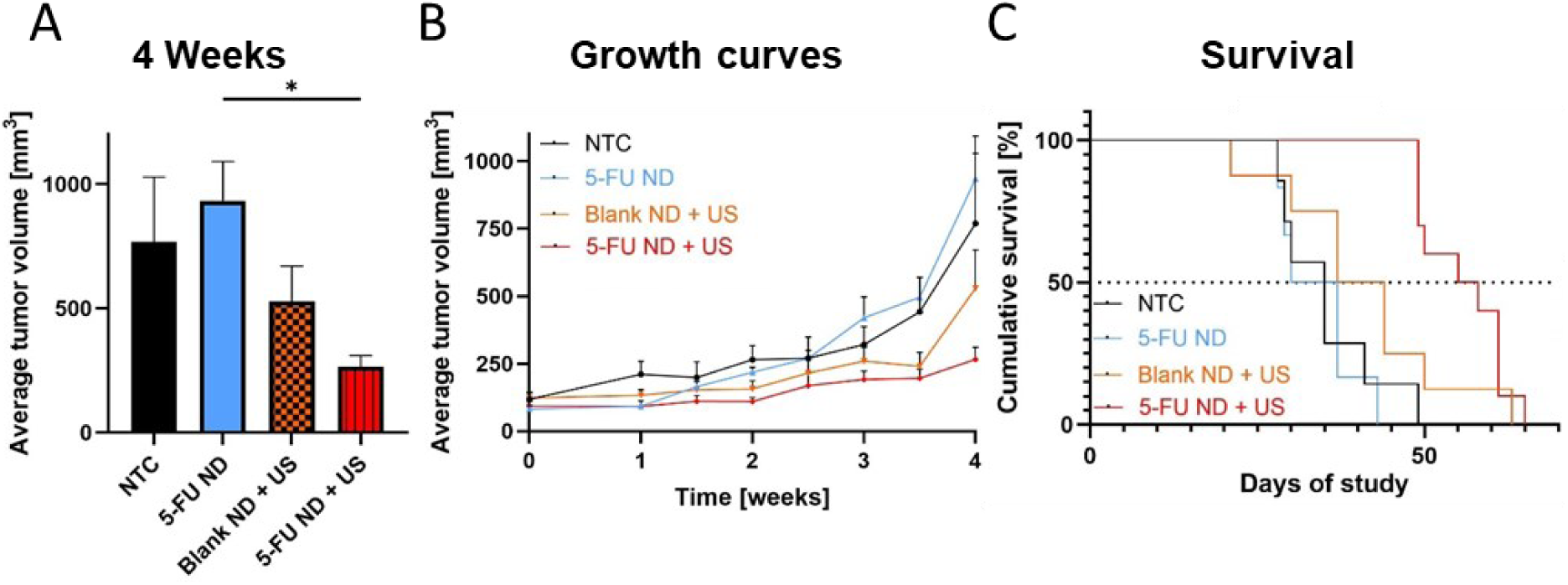
Therapeutic efficacy of ND-mediated mechanotherapy and chemotherapy uncaging in vivo. (A) Average tumor volume measured in week 4 for the different treatments. (B) Tumor growth curves from day 1 (treatment initiation) to week 4 for NTC, 5-FU ND, blank ND + US, and 5-FU ND + US groups. The 5-FU ND + US group shows sustained tumor suppression over the treatment period. (C) Kaplan-Meier survival curves displaying cumulative survival percentage over the experimental duration. The combined mechanochemical treatment (5-FU ND + US) confers a significant survival advantage compared to all other groups. Data represent mean ± SD. Statistical significance was determined by one-way ANOVA with Tukey’s post-hoc test (*p < 0.05, **p < 0.01, ***p < 0.001, ****p < 0.0001).

These growth trends were reflected in survival outcomes (Figure 8c). Median survival was 32.5 days in the NTC group and 33.5 days in the 5-FU ND group (without US), while blank ND + US extended survival to 40.5 days. The longest survival was observed in the 5-FU ND + US group, with a median survival of 56.5 days, corresponding to a 73.8% increase relative to NTC, a 68.7% improvement over 5-FU ND alone, and a 39.5% increase compared to blank ND + US. Survival analysis confirmed significant differences between groups (log-rank test p = 0.0001), with additional support from the log-rank trend test (p < 0.0001) and the Gehan-Breslow-Wilcoxon test (p = 0.0003). Together, these results demonstrate that ultrasound activation of 5-FU NDs produces synergistic antitumor effects and significantly improves survival.

## Discussion

This work aims to establish a robust solid tumor therapy platform by bringing together three elements into a single nano therapeutic agent: microfluidic production of small and stable lipid NDs, core loading of therapeutic as a platform for drug delivery, and a dual-frequency US protocol that couples ADV with low-frequency cavitation and drug release. The microfluidic approach enabled precise control over ND size and concentration. Increasing the total flow rate produced smaller (Table 1) and more concentrated droplets, consistent with other microfluidic studies where faster mixing favor smaller droplet formation. Between the two PFC cores we tested, C_5_F_12_ and C_6_F_14_, C_5_F_12_ gave smaller droplets at a given flow rate but proved less stable over time and temperature, likely because its boiling point (29.5°C vs 56°C of C_6_F_14_) is close to room and body temperature. C_6_F_14_ NDs, in contrast, maintained a stable size over 60 days at 4°C (Figure 2d) and remained liquid and intact at 37°C unless we applied focused US (Figure 2h). Their more negative zeta potential (−29.8 ± 3.7 mV compared to −20.58 ± 2.95 mV) and lower size fluctuation (Figure 2d-f) further support the choice of C_6_F_14_ as the preferred core for in vivo work.

The acoustic experiments clarify how these physical properties translate into functional behavior. In tissue-mimicking phantoms (Figure 2g), C_6_F_14_ NDs remained inert at room temperature and at 37°C for MI values below 1.0, compared to less stable C_5_F_12_ NDs, then gradually activated as the MI increased. We saw the first signs of vaporization at MI around 1.58 and full ADV with strong contrast enhancement at MI of 1.9 or higher (Figure 2h,i). This threshold-dependent activation is critical for clinical translation, as it ensures predictable vaporization only within targeted treatment zones while remaining safe during diagnostic imaging.

The core-loading strategy for 5-FU and BODIPY dyes is another important aspect. Rather than attaching the drug or dye to the shell, we dissolve both in the C_6_F_14_ phase before droplet formation. Many previously described PFC NDs carry drugs in the shell or in appended polymer layers, which can compromise stability and allow slow leakage over time. In our case, fluorination of the payloads allows them to stay inside the PFC core until the liquid-to-gas phase transition is triggered by US. The in vitro release data support this mechanism: after dual-frequency activation, BODIPY fluorescence in the supernatant increased about 2.8-fold (Figure 4c), and HPLC quantification showed a roughly 4-fold rise in 5-FU concentration (Figure 4f). These are substantial changes and indicate that a large fraction of the loaded drug is released during or immediately after ADV and cavitation. Multispectral IVIS analysis further demonstrated that the BODIPY-loaded NDs are well suited for optical tracking. Spectral characterization of the different BODIPY ND formulations enabled precise determination of their excitation and emission properties and confirmed clean separation of their fluorescence signatures, supporting multiplexed detection and distinct tracking of individual ND populations in heterogeneous biological environments (Figure 3d). This capability is important for future applications in which multiple ND subtypes (e.g., diagnostic vs therapeutic, or different drug payloads) may need to be monitored simultaneously in vivo.

The biological data show how these physical and acoustic properties translate into therapy. In vitro studies with 4T1 breast cancer cells were essential for evaluating blank ND cytotoxicity prior to US-mediated detonation. ND concentrations ranging from 25-200 µL were tested, and concentrations above 50 µL (1.26 x 10^10^ NDs) resulted in reduced cell viability (Figure 5c). This concentration-dependent toxicity may be attributed to the phospholipid shell components. At high phospholipid concentrations, previous studies have reported in vitro cytotoxicity due to membrane disruption or metabolic interference^62^. This likely accounts for the minor toxicity observed at elevated ND concentrations in vitro. It is important to note, however, that cells in culture are more sensitive to such effects than they would be in the complex and the supporting biological environment, where biological fluids, and clearance mechanisms may mitigate direct nanoparticle-cell interactions. Then, blank NDs at the chosen concentration (1.26 x 10^10^ NDs), and US alone at the parameters we used caused little damage (∼2.7% reduction). When we combined NDs with dual-frequency US, however, cell viability dropped to about 15% (∼85% reduction). This suggests that the two-step US protocol, first inducing ADV with the imaging transducer (keeping MI below 1.9 while applying acoustic pressure over 2 MPa), followed by stronger inertial cavitation with the low-frequency transducer, produces meaningful mechanical disruption even without drug present, consistent with reports of mechanical tissue fractionation in histotripsy and related techniques. The low-frequency transducer was selected over a high-frequency one to promote inertial cavitation of the gas bubbles generated during ADV, as lower frequencies (e.g., hundreds of kHz) lower the inertial cavitation threshold compared to MHz frequencies, enabling more efficient bubble collapse, microjet formation, and mechanical disruption at clinically feasible acoustic pressures. When 5-FU was encapsulated in the NDs and US was applied in the presence of tumor cells, viability fell further to around 7%, much lower than what we saw with passive 5-FU ND exposure (Figure 5e). This pattern supports a synergistic effect, a combination of mechanical damage from the cavitation and controlled chemotherapy effect.

In vivo, we see a consistent picture. BDPfr NDs showed higher and more persistent tumor signal than free BDPfr dye, both in IVIS in vivo and ex vivo imaging (Figure 6). At 24 hours, tumor fluorescence in ND-treated animals was 2.87-fold higher than in those given free dye, whereas free dye signals had largely returned to baseline in most tissues (Figure 6c,e). This indicates that the ND formulation improves both delivery to, and retention in, the tumor, as expected for a nanoscale carrier operating under the EPR effect. It is well-established that endothelial permeability increases in pathological states such as tumors and infarcted areas, enabling extravasation of large molecules and particles (10-500 nm, e.g., micelles, liposomes) into the interstitial space via EPR^13–15,63^. Nanoparticles with hydrodynamic diameters of 100-400 nm are considered optimal for passive tumor targeting through EPR, balancing vascular extravasation with retention while exceeding the renal clearance threshold (>40 kDa or ∼10 nm diameter), as glomerular sieving restricts first-pass kidney elimination of smaller species. In this context, the strong BDPfr ND signal persisting in tumors 24 h post injection reflects robust tumor targeting, ND stability in circulation, and sustained intratumoral retention, while the relatively low off-target signal in most organs supports a favorable biodistribution profile. Confocal microscopy (Figure 6f) goes a step further by showing that these NDs not only accumulate in the tumor but also reach deep tissue regions and intracellular compartments, indicating that the macroscopic retention measured by IVIS corresponds to genuine intra-tumoral distribution and cellular uptake rather than surface-restricted binding. To isolate mechanotherapy effects prior to combined treatment evaluation, blank NDs were tested with dual-frequency US (Figure 7a) and H&E staining of tumor sections (Figure 7b,d) revealed cavitation-induced lesions in the NDs + dual-frequency group compared to other groups (mean lesion areas: 1.55 ± 0.92% for untreated, 3.42 ± 1.39% for US only, 16.32 ± 5.02% for NDs + dual-frequency group).

The survival and tumor growth data confirm that these physical and imaging advantages have real therapeutic consequences (Figure 8). Weekly treatment with 5-FU NDs plus dual-frequency extended median survival from about 32.5 days in controls to 56.5 days (73.8% increase). Blank NDs with US gave an intermediate benefit, consistent with a contribution from purely mechanical damage, but the best outcome came from the combined chemomechanical therapy, demonstrating that our platform combines effective tumor targeting with potent mechanochemical therapy, demonstrating that our platform combines effective tumor targeting, deep tissue penetration, and potent mechanochemical therapy within a single US-controllable carrier.

The platform’s clinical translation potential is supported by several factors: microfluidic synthesis is compatible with GMP-scale production, PFCs have decades of clinical safety data, all acoustic parameters (MI ≤1.9) comply with FDA guidelines, and the dual-frequency system was assembled using off-the-shelf transducers. The ability to achieve therapeutic efficacy with reduced systemic drug exposure addresses a major limitation of conventional chemotherapy, and the demonstrated capacity for multispectral optical tracking and robust tumor-selective retention and intracellular delivery further supports the use of these core-loaded NDs as a modular theranostic platform.

In terms of future directions, while this study demonstrated efficacy in a breast cancer model, the platform’s versatility could extend to other indications and additional cancer types. NDs have been shown to enable blood brain barrier (BBB) opening via acoustic activation ^64,65^. It would therefore be valuable to explore this platform for targeted drug delivery in brain cancer and other neurodegenerative diseases, such as Alzheimer’s and Parkinson’s disease. In addition, the mechanochemical mechanism demonstrated here could be adapted for targeted therapy in inflammatory diseases, potentially reducing systemic immunosuppression associated with current treatments. Moreover, the tumor fractionation achieved by this approach may enhance immune infiltration and activation, suggesting that combining this method with immunotherapy represents a promising future direction.

## Materials and Methods

### Chemicals and Cell Culture

1,2-Distearoyl-sn-glycero-3-phosphocholine (DSPC, 850365P), and 1,2-distearoyl-*sn*-glycero-3-phosphoethanolamine-*N*-[meth-oxy(polyethyleneglycol)-2000] (ammonium salt) (DSPE-PEG2K, 880120P) were purchased from Avanti Polar lipids (Alabaster, AL, USA). Ethanol absolute (cat. no. 000525052100) was provided by Bio-Lab. PBS(-/-) 1× (phosphate-buffered saline without calcium and magnesium) was purchased from Sartorius (Cat# 02-023-1A), PBS(+/+) 1× (phosphate-buffered saline with calcium and magnesium) was purchased from Sartorius (Cat# 02-020-1A), Glycerol anhydrous was purchased from Bio-Lab, Propylene Glycol from Sigma-Aldrich (1,2-propanediol, Cat# 398039-500, St. Louis, MO, USA), BODIPY 493/503 (BDPg, 99%, CAS Number: 121207-31-6), Wheat germ agglutinin (WGA) Alexa Fluor 488 (W11261, Invitrogen) and 5-Fluorouracil (F6627, CAS Number: 51-21-8) were purchased from Sigma-Aldrich (St. Louis, MO, USA), BDP TR-X-NHS ester (BDPr, 95%, CAS Number: 197306-80-2) and BDP® 630/650-X-NHS ester (BDPfr, 95%, CAS Number: 2213445-35-1) were purchased from Lumiprobe, Agarose powder was purchased from Thermo Fisher (A10752, Thermo Fisher Scientific, OR, USA), Dulbecco modified Eagle medium (DMEM, Biological Industries [BI] Ltd., Cat # 01-055-1A, Kibbutz Beit-Haemek, Israel), RPMI 1640 medium w/ L-Glutamine was provided by Biological Industries Israel Beit-Haemek Ltd. (Cat # 01-100-1A), TrypLE Express dissociation reagent (Gibco Corp, 12604-013, Grand Island, NY, USA), 4T1 cells were acquired from ATCC (CRL-2539) and Met-1 mouse breast carcinoma cells were a gift from Prof. Jeffrey Pollard, University of Edinburgh, Edinburgh, UK, and Prof. Neta Erez, Tel Aviv University, Tel Aviv, Israel. Cell cultures were maintained in an incubator at 37°C in a controlled atmosphere (5% CO_2_, 95% humidity) using the culture media required by the producer using T25 and T75 flasks. All chemicals were used without further purification.

### Microfluidic fabrication of NDs

The preparation of C_5_F_12_ and C_6_F_14_-core lipid-shell NDs was achieved using a two-step microfluidic fabrication process utilizing the NanoAssemblr platform (v 1.5, Precision NanoSystems Inc., Vancouver BC, Canada) or by using Herringbone chip (LTF-012.00-4264, Darwin Microfluidics, France). In microfluidic technology, aqueous and organic phases are positioned in distinct streams within microchannels and are mixed through controlled laminar flow, where diffusion across their interface enables precise and efficient blending without turbulence. The aqueous phase was prepared by combining PBS(-/-) (pH 7.4), glycerol, and propylene glycol in an 8:1:1 v/v ratio. The mixture was then subjected to vortex mixing for 2 minutes. Following this, the mixture was sonicated in a water bath at 62°C for 15 minutes until it became homogeneous. The organic phase consisted of DSPC and DSPE-PEG2K (9:1 molar ratio) dissolved in ethanol (1.2 mg/mL lipid concentration), sonicated at 62°C until clear. The aqueous and organic solutions were maintained at 5±3°C before synthesis. PFC (C_5_F_12_ or C_6_F_14_) was added to the cold organic solution at 10 μL/mL ethanol just before use in the microfluidic system. Then, the two cold mixtures were vigorously stirred for 2 minutes using a vortex mixer until a homogeneous solution was achieved. Each mixture was then transferred into individual syringes and connected to the inlets of a microfluidic chip for further processing. To prepare encapsulated NDs, the process follows the same protocol as for regular NDs, with an additional step to incorporate the payload. For dye-loaded NDs (BDPg, BDPr, and BDPfr), the fluorescent dye is dissolved in the PFC (either C_5_F_12_ or C_6_F_14_) at a concentration of 0.5 mg/mL prior to its addition to the ethanol solution, alternatively the fluorescent dye can be dissolved in the pre-cooled ethanol solution containing the lipids and PFC at 0.05 mg/mL. Similarly, for drug-loaded NDs, the therapeutic chemotherapy (5-Fluorouracil) is dissolved in the pre-cooled ethanol solution containing the lipids and PFC at 0.1 mg/mL.

Two types of microfluidic-generated NDs were compared. Nanoassembly NDs synthesis was performed using the NanoAssemblr chip (Precision Nanosystems, 1146-039) with a total flow rate (TFR) of 12 mL/min and a flow rate ratio (FRR) of 3:1 (aqueous:organic), using the microfluidic mixing device Nanoassemblr. For the formation of other NDs under lower TFRs (100-2750 uL/min, FRR 3:1), a microfluidic system that included a Herringbone Mixer Glass Chip, and two NE-300 InfusionONE Syringe Pumps (NE-300-ES, Darwin Microfluidics, France) was used.

Following microfluidic synthesis, the suspensions were purified by dialysis using a Pur-A-Lyzer™ Maxi Dialysis Kit (12-14 kDa MWCO, 3 mL capacity) under gentle stirring at 4°C. For standard preparations, dialysis was performed against phosphate-buffered saline (PBS-/-, pH 7.4) for 24 h with buffer replacements at 3 h and 6 h (1:200 v/v sample-to-PBS ratio) to remove residual ethanol, unencapsulated phospholipids, fluorescent molecules, and free therapeutic drugs. For TEM and E-SEM samples, deionized water was substituted as the dialysis medium (22 h total, with buffer changes at 3 h and 6 h). Post-dialysis, all samples were stored at 4°C. Formulation Example: The aqueous phase comprised PBS (32 mL), glycerol (4 mL), and propylene glycol (4 mL), totaling 40 mL. The organic phase consisted of DSPC (22.7 mg), DSPE-PEG2k (2.5 mg), EtOH (10 mL), and C_5_F_12_ (100 μL). Microfluidic parameters included for example organic flow rates of 25, 75, and 125 μL/min, with corresponding aqueous flow rates of 75, 225, and 375 μL/min, resulting in FRR of 3:1 and TFR of 100, 300, and 500 μL/min.

### NDs characterization

The size distribution and concentration of the NDs were characterized using Nanoparticle Tracking Analysis (NTA, NanoSight NS300, Malvern Panalytical, UK) with software version NTA 3.4 Build 3.4.4, Dynamic Light Scattering (DLS, Zetasizer Ultra, Malvern Panalytical, UK), and, for selected samples, Prometheus Panta system (NanoTemper Technologies, Germany). Zeta potential was measured using Zetasizer Ultra (DLS, Zetasizer Ultra, Malvern Panalytical, UK). The morphology of the NDs was further examined using Transmission Electron Microscopy (TEM, JEM 1400plus) and Environmental Scanning Electron Microscopy (E-SEM) was performed using the Quanta FEG 650 (Thermo Fisher Scientific, USA) at an accelerating voltage of 30 kV, equipped with a Solid-State Detector (SSD) (Thermo Fisher Scientific, USA). The physical stability of the NanoAssemblr-NDs in suspension was evaluated by monitoring their size distribution over time. To assess the stability of the NDs under storage conditions, the droplets were stored at 4°C for up to one month (with blank C_5_F_12_ and C_6_F_14_ NDs samples extended to two months for additional evaluation). The ND suspensions were 10-fold diluted with deionized water and analyzed using a Zetasizer Ultra DLS instrument. To evaluate the stability of C_5_F_12_ and C_6_F_14_ NDs under physiological conditions, the suspensions were maintained at 25°C, 37°C, and 42°C in a Panta DLS device for up to one hour. Size distribution measurements were taken at regular intervals to monitor changes in droplet stability under these conditions.

### Ultrasound imaging of ND activation thresholds

ND activation was monitored using an L7-4 linear array transducer (Philips, ATL, 5.5 MHz center frequency, 128 elements) mounted on a programmable Vantage 256 ultrasound system (Verasonics Inc., Redmond, WA, USA). The same transducer delivered focused acoustic pressures and acquired B-mode images pre/post-activation. All experiments were performed in triplicate.

Tissue-mimicking agarose phantoms (1.5% agarose in deionized water) contained cylindrical inclusions (3D-printed mold: 65 × 25 × 20 mm outer dimensions, 15 mm tall × 8 mm diameter central rod) for ND activation and in vitro experiments, filled with ND suspension (10 µL NDs in 1 mL degassed PBS (-/-)). Phantoms were positioned with the ND inclusion at the transducer focal point. Experiments characterized acoustic vaporization thresholds and thermal stability at 25°C, 37°C, and 42°C using MI 1.5-2.1 for up to 2 hours. ND-to-MB phase transition was detected as echogenicity enhancement due to gas-core acoustic impedance mismatch, visualized in real-time B-mode imaging.

### Dual-frequency focused ultrasound system

The US setup was adapted from our previous study^43,66^ and consisted of a dual-frequency system integrated within a water tank. A spherically focused, single-element therapeutic transducer (H149, Sonic Concepts, Bothell, WA, USA, 105 kHz center frequency, 65 mm focal length) was coaxially aligned with a 1D rotating imaging probe (IP104, Sonic Concepts, Bothell, WA, USA) positioned at its base. The therapeutic transducer was driven by a TPO-200 amplifier (Sonic Concepts) and connected through a matching network (Sonic Concept). The 128-element imaging probe (13.5 mm elevation × 28.2 mm aperture) was controlled by the Verasonics system and mounted on a motorized rotary stage (RTY-IP100, Sonic Concepts) capable of ±180° rotation, with motion parameters governed by custom MATLAB scripts.

Transducers calibration was performed using a 0.2 mm needle hydrophone (NH0200, Precision Acoustics, Dorchester, U.K.) and a 3D positioning system (Newport motion controller ESP 300). The full-width at half-maximum (FWHM) of the focal zones was characterized as 1.2 mm (x) × 2.6 mm (y) × 23 mm (lateral) for the imaging probe and 10 mm (lateral) × 40 mm (axial) for the therapeutic transducer. The PNP for both transducers were calibrated in situ. For NDs vaporization, the imaging probe delivered 2-cycle sinusoidal pulses (3.5 MHz, MI 0.84) with single-focus steering and rotation to cover the treatment volume. This probe also facilitated in vivo tumor imaging and positioning within the focal area.

### Fluorescent labeling and release characterization

Fluorescent NDs encapsulated fluorine-functionalized BODIPY derivatives in green (BDPg), red (BDPr), or far red (BDPfr). To assess release kinetics, BDPg NDs were evaluated using a SPARK multimode microplate reader (Tecan Group Ltd., Männedorf, Switzerland) in fluorescence mode. Experiments were performed in 96-well plates with data acquisition and analysis via SPARKControl software. US-triggered release was evaluated using a dual-frequency system incorporating a 3.5 MHz imaging transducer (MI 0.84) and a 105 kHz therapy transducer (MI 1.67) operating concurrently. Top supernatant solutions collected before and after US exposure, diluted 10-fold with DI and filtered through a 0.22-µm membrane filter, then the fluorescence intensity was measured to quantify the extent of fluorophore release by microplate reader and spectrophotometer, using a total measurement volume of 200 µL per well in a 96-well format. The change in fluorescence intensity between pre- and post-US treatments represented the release of fluorescent payload. Calibration was performed using free dye standards at concentrations ranging from 0.0 to 0.6 µM in Chloroform (0.0, 0.1, 0.143, 0.2, 0.3, 0.4, 0.5, and 0.6 µM). The excitation and emission wavelengths were 493 ± 5 nm and 513 ± 5 nm, respectively.

### 5-Fluorouracil (5-FU) chemotherapy encapsulation and release quantification

To validate the differential solubility of 5-FU in C_6_F_14_ versus aqueous media, 5-FU was dissolved in either C_6_F_14_ or DI water at a concentration of 0.4 mg/mL. Following vigorous mixing, solutions were filtered through a 0.22-µm membrane filter to remove undissolved particulates. 5-FU concentration can be estimated by UV-spectrophotometer^67^. Fluorescence intensity was evaluated using a Spark multimode microplate reader (Tecan Group Ltd., Männedorf, Switzerland). Samples (pure DI water, pure C_6_F_14_, 5-FU in DI, and 5-FU in C_6_F_14_) were loaded into UV-transparent 96-well plates while scanned wavelengths were within UV spectrum, and data were acquired using SparkControl software. A preliminary full-spectrum scan was performed to identify optimal peak wavelengths (excitation: 240-295 nm, emission: 310 ± 5 nm), followed by quantitative endpoint analysis using optimized parameters (excitation: 265 ± 20 nm, emission: 310 ± 20 nm). All measurements were performed in triplicate, and background signals from pure solvents were subtracted during analysis.

Quantification of 5-FU release from 5-FU NDs (1.18 × 10^11^ ND/mL) was conducted using high-performance liquid chromatography (HPLC, Agilent 1260 Infinity Series) equipped with a photodiode array UV-vis detector. Supernatants collected before and after US exposure (under the same US parameters as fluorescence release experiment with dual-frequency set-up) were diluted 100-fold with deionized water, prefiltered through 0.22 µm membranes, and transferred to 2 mL HPLC vials prior to injection. A 50 µL sample was injected into a Fast-Poroshell 120 EC-C18 column (100 mm × 4.6 mm, 4 µm, Agilent Technologies, USA) maintained at 30°C. The mobile phase was an isocratic mixture of 4% HPLC-grade methanol (Bio-Lab Ltd., Israel) and 96% deionized water containing 0.1% acetic acid (Merck KGaA, Germany), at a flow rate of 0.8 mL min^-1^. Detection was performed at 254 nm with a retention time of 2.2 min, consistent with established analytical protocols^68,69^. Calibration was based on free 5-FU standards ranging from 0.005 to 0.1 ppm.

### Transmission Electron Microscopy

Transmission Electron Microscopy (TEM) was conducted using the JEM 1400plus transmission electron microscope (Jeol, Japan). Samples were adsorbed onto Formvar/carbon-coated grids and stained with 2% aqueous uranyl acetate prior to imaging. Images were captured using the SIS Megaview III camera and iTEM imaging platform (Olympus, Japan). For sample preparation (negative staining approach), 10 µL samples were diluted with DI to 1 mL, then 10 µL solutions were adsorbed on formvar/ carbon coated grids and stained with 2% aqueous uranyl acetate. Afterward, the grid was dried in air for 1 hour at room temperature and observed using TEM.

### Environmental Scanning Electron Microscope

The microstructure and morphology of the samples were characterized using an Environmental Scanning Electron Microscope (E-SEM, ThermoFisher, Quanta 200 FEG). The system is equipped with a Schottky field-emission gun electron source, operating at 30 kV accelerating voltage and beam currents up to 100 nA, offers three vacuum modes - high vacuum, low vacuum, and environmental/wet modes, while we used the environmental mode for our samples to maintain the integrity of the liquid phase during imaging. For liquid sample preparation, a 10 µL aliquot of the solution was carefully adsorbed onto a formvar/ carbon coated grid. The grid was allowed to dry under ambient conditions for 2 minutes to ensure proper adsorption of the sample.

### In vitro evaluation of NDs in 4T1 breast cancer cells

In vitro studies were conducted using 4T1 cells, a metastatic triple negative murine breast carcinoma cell line (purchased from ATCC), to evaluate the therapeutic potential of C_6_F_14_ ND-mediated mechanotherapy. The cells were cultured in RPMI 1640 Medium (With ʟ-glutamine, supplemented with 10% v/v fetal bovine serum, 1% v/v penicillin-streptomycin) at 37°C in a humidified 5% CO_2_ incubator. On the day of the experiment, cells (≈85% confluency) were dissociated with TrypLE Express (Gibco Corp, 12604-013, Grand Island, NY, USA) and resuspended at 1 × 10^6^ cells/mL in degassed PBS containing calcium and magnesium (PBS+/+).

Prior to US experiments, we conducted a concentration optimization study to determine the non-cytotoxic range of C_6_F_14_ NDs (blank NDs) for 4T1 breast cancer cells. Cells were seeded in 6-well plates at a density of 2×10^5^ cells per well in 2mL RPMI 1640 Medium (With ʟ-glutamine, supplemented with 10% v/v fetal bovine serum, 2.5% v/v penicillin-streptomycin) and exposed to increasing volumes of NDs (25-200 μL of 2.52×10^11^ particles/mL suspension, with three replicates each), with medium replacement at 48 hours to maintain nutrient availability and remove non-adherent cells. After 72 hours of exposure, cells were collected in TrypLE Express, and viability was quantified via direct cell counting and normalized to non-treated controls (NTC) to account for baseline proliferation. This screening identified the optimal NDs volume that maintained >90% viability while ensuring high ND concentration for therapeutic experiments. All treatments were analyzed in triplicate.

For the US-mediated therapy experiments, 4T1 cells were prepared at a concentration of 1×10^6^ cells/mL in PBS containing calcium and magnesium. Aliquots of 200 μL (containing 2×10^5^ cells) were distributed into 0.5 mL Eppendorf tubes and divided into six experimental groups with three replicates each. The treatment groups included NTC, US alone (combined 105 kHz and 3.5 MHz transducers), NDs alone at the predetermined optimal concentration (1.26 × 10^10^ NDs), and three combination groups pairing NDs with either imaging US transducer, therapeutic US transducer, or dual-frequency system containing both. Hydrophone calibration inside the 0.5 mL Eppendorf tubes showed a 30% acoustic PNP attenuation, consistent with literature findings for similar setups^43^. Each treated sample was positioned at the focal point of our custom US system, featuring a rotating imaging transducer (3.5 MHz, MI 1.67) and a therapeutic transducer (105 kHz, MI 0.84) each operating for 120 seconds. The therapeutic transducer delivered pulsed waves with 0.5 ms burst length and 30 ms period. After treatment, cells were transferred to six-well tissue culture dishes containing RMPI 1640 medium supplemented with 2.5% v/v penicillin-streptomycin and cultured at 37°C in a humidified 5% CO_2_ incubator for 72 h, with medium replacement at 48 hours. Live cells were counted after TrypLE Express dissociation. All experiments were performed in triplicate.

### Combined mechanical-chemotherapeutic efficacy of 5-FU loaded NDs

In vitro studies of 5-FU NDs were conducted using breast cancer cells (4T1 with stable lentiviral expression of the ZsGreen1 protein for confocal microscopy, or regular 4T1 cells) to evaluate the therapeutic potential of US-mediated mechanotherapy and drug delivery of 5-FU. The cells were cultured in RPMI 1640 Medium (With ʟ-glutamine, supplemented with 10% v/v fetal bovine serum, 1% v/v penicillin-streptomycin) at 37°C in a humidified 5% CO_2_ incubator. On the day of the experiment, cells (≈85% confluency) were dissociated with TrypLE Express (Gibco Corp, 12604-013, Grand Island, NY, USA) and resuspended at 1 × 10^6^ cells/mL in degassed PBS containing calcium and magnesium (PBS+/+).

For the experiments, 4T1 cells were prepared at a concentration of 1×10^6^ cells/mL in PBS containing calcium and magnesium. Aliquots of 200 μL (containing 2×10^5^ cells) were distributed into 0.5 mL Eppendorf tubes and divided into experimental groups with three replicates each. NDs (blank NDs or 5-FU NDs) at the predetermined optimal concentration from the previous experiment (1.26 × 10^10^ NDs). For the first experiment, containing regular 4T1 cells the groups included non-treated controls (group 1), blank NDs without US (group 2), 5-FU NDs without US (group 3), blank NDs + dual-frequency system (group 4), and 5-FU NDs + dual-frequency system (group 5). Each treated sample was positioned at the focal point of our custom US system, featuring a rotating imaging transducer (3.5 MHz, MI 1.67) and a therapeutic transducer (105 kHz, MI 0.84) each operating for 120 seconds. The therapeutic transducer delivered pulsed waves with 0.5 ms burst length and 30 ms period, ensuring synchronized ND vaporization (imaging), cavitation and drug release (therapeutic) effects. After treatment, cells were transferred to six-well tissue culture dishes containing RMPI 1640 medium supplemented with 2.5% v/v penicillin-streptomycin and cultured at 37°C in a humidified 5% CO_2_ incubator for 72 h, with medium replacement at 48 hours. Live cells were counted after TrypLE Express dissociation.

For the second experiment, GFP-expressing 4T1 breast cancer cells (4T1 with stable lentiviral expression of the ZsGreen1 protein) were used, divided into four groups: NTC (group 1), 5-FU NDs (1.26 × 10^10^ NDs) co-incubated with 4T1 cells without US exposure to assess passive drug release (group 2), 5-FU NDs (1.26 × 10^10^ NDs) exposed to dual-frequency US in Eppendorf tubes without cells, followed by addition of the resulting chemotherapy solution to 4T1 cell culture to isolate the chemotherapy effect (group 3), and 5-FU NDs (1.26 × 10^10^ NDs) combined with 4T1 cells subjected to simultaneous dual-frequency US application, representing mechanotherapy plus drug release (group 4). Each treated sample was positioned at the focal point of a custom US system comprising a rotating imaging transducer (3.5 MHz, MI 1.67) and a therapeutic transducer (105 kHz, MI 0.84), each operating for 120 seconds. The therapeutic transducer delivered pulsed waves with a 0.5 ms burst length and 30 ms period. Post-treatment, cells were transferred into six-well tissue culture plates containing RPMI 1640 medium supplemented with 2.5% penicillin–streptomycin and incubated at 37°C in a humidified atmosphere with 5% CO_2_ for 72 hours, with medium replaced at 48 hours.

Prior to confocal microscopy, fresh medium was supplied, and imaging was performed on an Olympus FV3000-IX83 system (Olympus, Japan) equipped with a UPLXAPO 20× objective (NA 0.7). GFP fluorescence was excited using a 488 nm laser (1.75% transmissivity at 100% power) and detected through the Alexa Fluor 488 channel. Paired brightfield and fluorescence images (512 × 512 pixels, 12-bit depth) were captured at a pixel size of 1.24 µm (0.6215 µm/pixel with 2× digital zoom) using a Galvano scanner in line sequential mode at 2.114 µs/pixel. Untreated GFP-expressing 4T1 cells served as imaging controls. All image processing was performed using ImageJ (NIH).

### Fluorescence characterization of NDs by multispectral IVIS imaging

The fluorescence profiles of BDPg ND, BDPr ND, and BDPfr ND were characterized using an IVIS imaging system. Five Eppendorf tubes were arranged (left to right: PBS control, BDPg ND, BDPr ND, BDPfr ND, and air blank) and imaged across 15 excitation/emission filter pairs spanning visible to near-infrared wavelengths (480-740 nm excitation steps of 20 nm, 620/710/790 nm emission). This multispectral approach enabled resolution of ND-specific fluorescence signals while accounting for potential background autofluorescence. All acquisitions used identical instrument settings to permit direct comparison between formulations.

### Mice breast cancer model and treatment protocol

All animal procedures were approved by the Institutional Animal Care and Use Committee (IACUC) of Tel Aviv University and were carried out in compliance with institutional guidelines for care and use of animal models (approval TAU-MD-IL-2506-132-4). MET-1^70^ mouse breast carcinoma cells were injected into 8 to 12 weeks old female FVB/NHanHsd mice (Envigo, Jerusalem, Israel). The cells were cultured at 37°C in a humidified 5% CO_2_ incubator in Dulbecco’s modified Eagle medium (DMEM, high glucose, supplemented with 10% v/v fetal bovine serum, 1% v/v penicillin-streptomycin and 0.11 g L^-1^ sodium pyruvate) until ≈85% confluency, then dissociated with TrypLE Express and resuspended in PBS+/+. During the day of the injection, Met-1 cells were collected with TrypLE Express dissociation reagent to a final concentration of 1 × 10^6^ cells in 25 μL of PBS+/+. The cells were subcutaneously injected into #4 inguinal mammary fat pads to establish tumors. The tumor size was recorded every 4 days until they reached approximately a volume of 80-100 mm^3^, unless otherwise specified for individual experiments. Mice were anesthetized with isoflurane (Piramal), the fur in the tumor area was shaved and depilated (Veet hair removal cream) and positioned within the US transducer focal zone for treatment.

### In vivo imaging and pharmacokinetic analysis

Biodistribution, tumor accumulation, and blood circulation profiles of BDPfr NDs versus free dye were characterized to establish clearance kinetics, organ distribution patterns, and tumor targeting efficiency prior to therapeutic studies

The mice were divided into 3 groups: NTC (n=3), free BDPfr (n=4) and BDPfr NDs (n=4) injection. 150 uL of diluted BDPfr NDs (150 μL total volume: 50 μL ND solution + 100 μL PBS(-/-)) or free BDPfr (150 μL at equivalent dye concentration (0.2 µM BDPfr) in 5% ethanol/PBS) were injected retro orbitally when the tumor volume reached an average of 1000 mm^3^, the mice were anesthetized using isoflurane. Fluorescence levels were assessed and analyzed with 640 nm excitation and 710 nm emission for whole mice at different time points during 24 h and different organs (tumor, liver, spleen, kidneys and brain) ex-vivo 24 h post injection using IVIS SpectrumCT (Caliper IVIS Lumina Series III) in vivo imaging system and Living Image 4.8.2 software (PerkinElmer, USA) using background subtraction and radiant efficiency quantification ([photons/s/cm²/sr]/[µW/cm²]).

In a separate pharmacokinetic experiment, the blood biodistribution and circulation time of BDPfr NDs were evaluated. Animals were assigned to either NTC (n=3) or BDPfr ND (n=3) groups. Once tumors reached a volume of approximately 1000 mm^3^, BDPfr NDs were administered via retro-orbital injection under isoflurane anesthesia (Piramal, Israel). The BDPfr ND group received 150 uL per 20 g mouse (50 uL ND solution + 100 uL PBS(-/-)), while the NTC group provided baseline fluorescence measurements. Serial blood samples were collected via cheek pouch puncture at 10 min, 3 h, and 24 h post-injection following a staggered sampling schedule adapted from established protocols to minimize animal stress^71^. Blood was immediately transferred into 96-well plates, and fluorescence was quantified ex vivo using an IVIS Spectrum/SpectrumCT system (Caliper IVIS Lumina Series III, PerkinElmer, USA) with 640 nm excitation and 710 nm emission. Radiant efficiency ([photons/s/cm²/sr]/[µW/cm²]) was calculated by defining regions of interest (ROIs) in Living Image 4.8.2 software (PerkinElmer), enabling comparative pharmacokinetic analysis between BDPfr ND treated and NTC groups.

### Tissue harvesting and histology protocol

For ex vivo therapeutic assessment, tumors were harvested. In addition, after the ex vivo biodistribution study using IVIS, tumors and internal organs (spleen, liver, kidneys, and brain) were harvested. Tumors and internal organs were frozen using liquid nitrogen and methyl butane and processed for embedding in O.C.T Compound (ThermoFisher Scientific, USA), and then stored at −80°C. Frozen samples were subsequently cryo-sectioned into 12 µm thick slices in a −20°C cryostat microtome (CM1950, Leica Biosystems), and processed via two parallel pathways: (1) for histological evaluation of US treatment, sections were stained with hematoxylin (Leica 3801542) and eosin (Leica 3801602) (H&E) following standardized protocols^72^ and stored at −80°C, (2) for fluorescent NDs biodistribution analysis, sections were mounted on glass slides and treated with WGA Alexa Fluor 488, washed with PBS and then treated with three drops of DAPI containing mounting medium (Fluoroshield, ab104139, Abcam), then coverslipped and allowed to develop for 15 minutes at room temperature in light protected conditions and stored at −80°C before further analysis.

### In vivo mechanotherapy in a breast cancer mouse model

Blank ND-mediated mechanical tissue fractionation was evaluated using the dual-frequency US system prior to combined therapy studies. Mice were randomized into four groups (n=3/group): (1) NTC, (2) NDs + imaging US (3.5 MHz, MI 1.84), (3) NDs + therapeutic US (105 kHz, MI 0.91), and (4) NDs + combined dual-frequency US. Anesthetized mice received retro-orbital injection of blank NDs (110 µL) and were positioned atop the water tank using an agarose spacer to place tumors at the dual-frequency focal zone after 10 min circulation. US was administered for 6 minutes (2-minute sonications per plane across three z-axis planes spaced by 2.5 mm to cover the tumor volume, given the imaging transducer’s 2.4 mm focal spot). Combined transducer groups received simultaneous imaging/therapeutic US. Mice were sacrificed at 24 h post-treatment, tumors were frozen in liquid nitrogen-cooled methylbutane (−80°C storage) and sectioned (12 μm, Leica CM1950 cryostat at −20°C) for H&E staining to assess tissue fractionation.

### Microscopy imaging and quantitative analysis

Following thawing to room temperature, tissue slides sections were imaged using the Echo Revolution microscope (Echo, San Diego, USA). Scans were performed at 4× and 20× optical magnification to assess both tumor fractionation morphology in H&E-stained sections (using the brightfield channel) and organ-specific NDs extravasation patterns in fluorescent samples. For fluorescence imaging, optimized excitation wavelengths were applied: 365 nm (DAPI), 410 nm (GFP), 550 nm (TRITC), and 690 nm (Cy5), allowing simultaneous visualization of cellular architecture and NDs distribution. This imaging protocol facilitated correlative analysis of therapeutic effects at tissue level and drug delivery efficiency at the cellular scale. For comparison and quantification of the resulting lesions, post-processing of the scanned images was performed in ImageJ. Each image was converted to a binary image, where the lesion areas were turned black, and non-lesion areas were white. The lesion area percentage was calculated using the following equation (1):

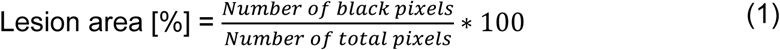

To assess intra-tumoral distribution of BDPfr NDs, tumors were harvested 24 h post injection and processed for embedding in O.C.T Compound (ThermoFisher Scientific, USA). The O.C.T blocks were cut into 12 μm sections, which were made serially through multiple regions of each tumor. Slides then were examined and imaged by Echo Revolution microscope (Echo, San Diego, USA) Then tumor slides randomly chosen and underwent WGA Alexa Fluor 488 (cell membrane), PBS washing, then DAPI immunomounting for cell nuclei as mentioned in tissue harvesting protocol section. Dried slides were examined and imaged by a TCS SP8 multiphoton confocal microscope (Leica, USA).

### Survival study

The long-term therapeutic effect of combined mechanotherapy and chemotherapy release was evaluated in bearing MET-1 breast carcinomas, with each mouse having a single tumor. Four groups were included in this study: 1) NTC group, 2) a group that received 5-FU ND treatment alone without US (5-FU ND), 3) Blank ND-mediated histotripsy (Blank ND + US) and 4) 5-FU ND-mediated histotripsy (5-FU ND + US) (n = 8 mice per group). US treatment was done with a dual-frequency system (3.5 MHz, MI 1.84 and 105 kHz, MI 0.91). Treatments were repeated weekly, and tumor growth was monitored twice a week. The mice were sacrificed when the tumor volume reached 1000 mm^3^.

In a separate endpoint imaging experiment, a subset of mice from the NTC, Blank ND + US, and 5-FU ND + US groups received an injection of BDPfr NDs when tumor volume reached 1,000 mm^3^. Tumors and organs were harvested and scanned ex vivo by IVIS 4 h post-injection. The imaging cohort consisted of four groups (n = 3 per group): control (no BDPfr NDs), BDPfr NDs only, BDPfr NDs after Blank ND + US treatment, and BDPfr NDs after 5-FU ND + US treatment.

## Statistical analysis

Prism 10.1.2 (GraphPad Software) was employed for statistical analysis. In experiments involving multiple groups, differences among multiple populations and subpopulations were assessed using One-Way ANOVA with Tukey’s multiple comparisons. The value of p ≤ 0.05 was considered statistically significant. Differences are presented on graphs in the following abbreviations: blank. for not significant, * for p ≤ 0.05, ** for p ≤ 0.01, *** for p ≤ 0.001, and **** for p ≤ 0.0001.

## Author Contributions

T.B designed and performed the research, analyzed the data, and wrote the manuscript. M.B, and M.G assisted with in-vivo experiments. D.P guided, advised and contributed to the manuscript. T.I. guided, advised, designed the research and contributed to the manuscript. All authors reviewed the manuscript.

## Declaration of generative AI use

During the preparation of this work the authors used ChatGPT (OpenAI) to assist with language editing and rephrasing certain sections of the manuscript. After using this tool, the authors reviewed and edited the content as needed and they take full responsibility for the content of the published article.

## Data Availability

The datasets generated during and/or analyzed during the current study will be made available upon request.

