## Supplementary information for "Acoustically activatable drug-loaded nanodroplets for mechanochemical therapy in solid tumors"

**This PDF file includes:**
Supplementary Tables
Table S1

Supplementary Figures
Fig. S1-S6

**Table S1 - Physical characteristics of PFC and other materials useful for making blank NDs and encapsulated NDs.**

| Compound | Molecular Weight (g/mol) | Boiling Point (°C) | Phase at RT | Solubility in Water | Excitation/ Emission Wavelengths (nm) |
|---|---|---|---|---|---|
| Perfluoropentane ($C_5F_{12}$) | 288.05 | 29.5 | Liquid | Insoluble | NA |
| Perfluorohexane ($C_6F_{14}$) | 338.06 | 56 | Liquid | Insoluble | NA |
| 5-Fluorouracil (5-FU) | 130.08 | Sublimes at 282-283 | Solid | Highly Soluble in organic solvents, low solubility in water. | Absorption: 265 |
| BODIPY 493/503 | 262.11 | NA | Solid | Soluble in organic solvents, Insoluble in water. | Ex: 493, Em: 503 |
| BDP TR-X-NHS ester | 634.46 | NA | Solid | Soluble in DMSO, DMF. Insoluble in water. | Ex: 589, Em: 616 |
| BDP® 630/650-X-NHS ester | 660.5 | NA | Solid | Soluble in DMSO, DMF. Insoluble in water. | Ex: 628, Em: 642 |

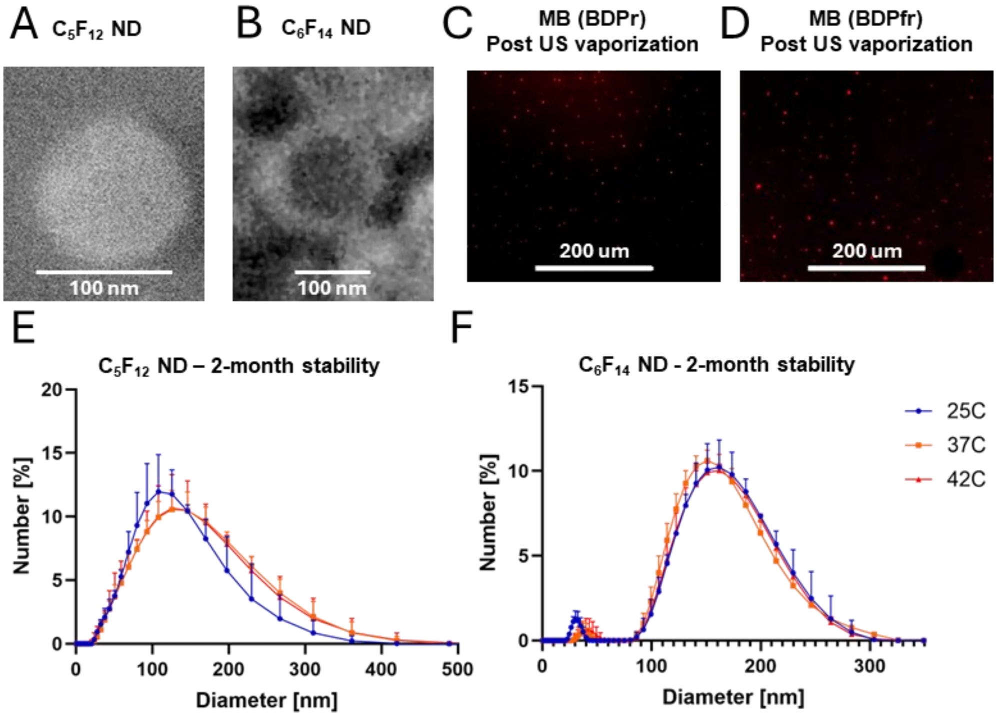


**Figure S1. Structural, and size stability of perfluorocarbon nanodroplets.** (A) ESEM image of $C_5F_{12}$ NanoAssemblr-formulated NDs (scale bar: 100 nm). (B) ESEM image of $C_6F_{14}$ NanoAssemblr-formulated NDs (scale bar: 100 nm). (C) Fluorescence microscopy of BDPr-loaded microbubbles generated by ultrasound vaporization of BDPr NDs (scale bar: 200 µm). (D) Fluorescence microscopy of BDPfr-loaded microbubbles generated by ultrasound vaporization of BDPfr NDs (scale bar: 200 µm). (E) Number-weighted DLS size distributions of $C_5F_{12}$ NanoAssemblr NDs after 2 months storage at 2-8 °C, measured at 25, 37, and 42 °C, showing a temperature-dependent shift of the distribution toward larger diameters at 37-42 °C compared with 25 °C, consistent with droplet expansion and/or microbubble formation. (F) Number-weighted DLS size distributions of $C_6F_{14}$ NanoAssemblr NDs after 2 months storage at 2-8 °C, measured at 25, 37, and 42 °C, demonstrating overlapping distributions and stable mean diameters across all temperatures.

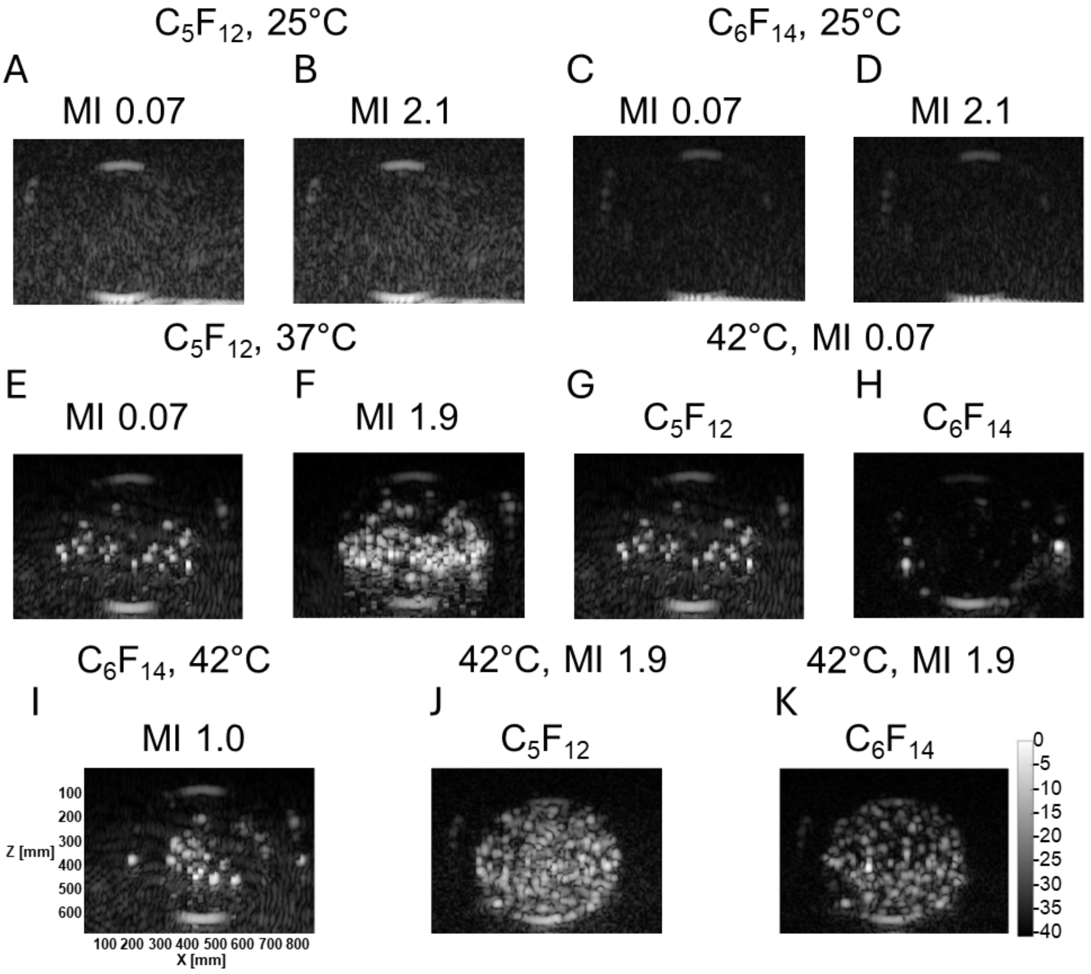


**Figure S2. Acoustic droplet vaporization of $C_5F_{12}$ and $C_6F_{14}$ nanodroplets at 12 mL/min flow rate under varying temperature and mechanical index.** Representative B-mode ultrasound images acquired from an agarose tissue-mimicking phantom containing ND suspensions using an L7-4 linear array transducer (5.5 MHz). At 25°C, neither $C_5F_{12}$ (a,b) nor $C_6F_{14}$ (c,d) NDs exhibited activation, even at high MI (2.1), confirming thermal stability at ambient temperature. At 37°C, $C_5F_{12}$ NDs showed no activation at low MI (0.07, e) but underwent pronounced vaporization at MI 1.9 (f), demonstrating pressure-dependent phase transition. $C_5F_{12}$ and $C_6F_{14}$ NDs at 42°C showed minimal activation at low MI (0.07, g,h) and $C_6F_{14}$ NDs showed partial activation at MI 1.0 (i), while both formulations exhibited enhanced activation at MI 1.9 (j,k), reflecting combined thermal and acoustic triggering. Scale bar is common to all subfigures.

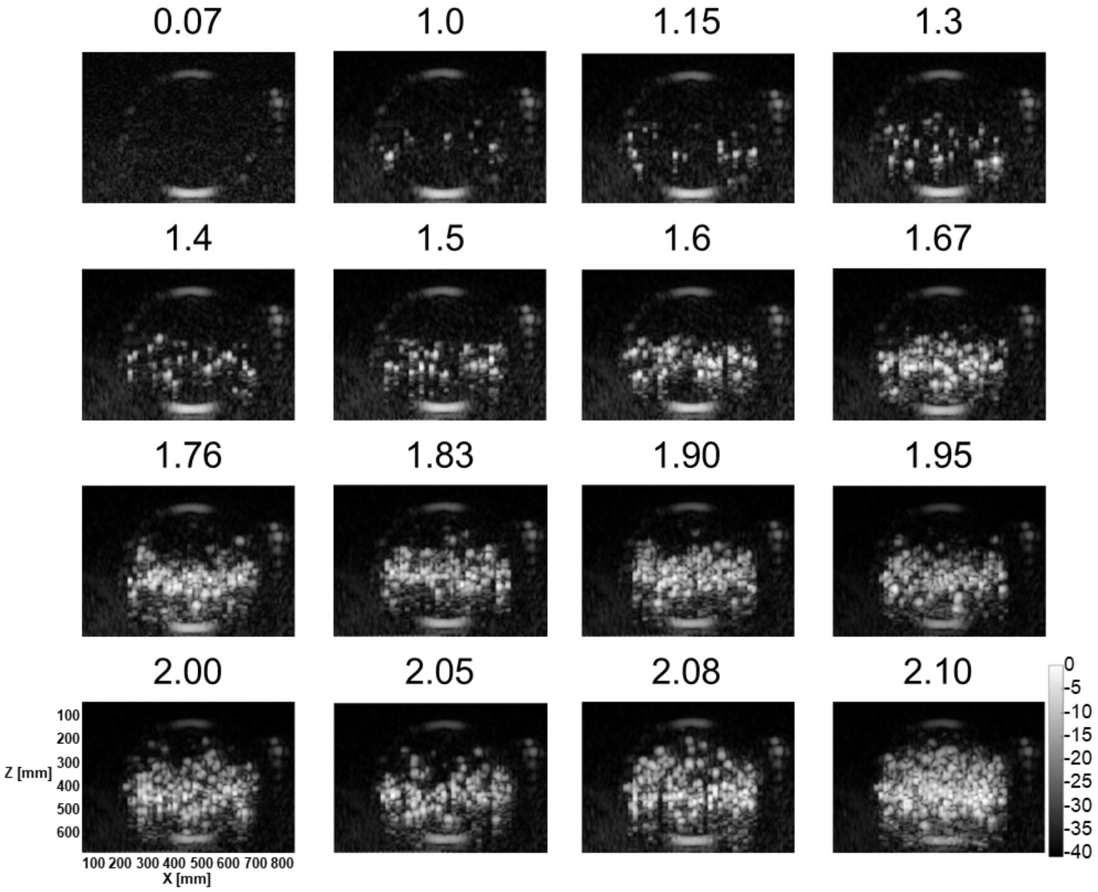


**Figure S3. Acoustic imaging of $C_6F_{14}$ NanoAssemblr nanodroplet vaporization at 37 °C under varying mechanical indices.**

Representative B-mode ultrasound images acquired from an agarose tissue-mimicking phantom containing a $C_6F_{14}$ ND suspension using an L7-4 linear array transducer (5.5 MHz). Images were captured at increasing mechanical indices (MI = 0.07-2.10). At low MI (0.07), no acoustic droplet vaporization (ADV) or microbubble formation is observed. Sparse microbubbles begin to appear at MI ≈ 1.0, with progressive vaporization and microbubble accumulation as MI increases (1. 0-2.1). Pronounced ADV occurs at MI ≥ 1.67, reaching maximal microbubble generation near the highest tested MI (2.1). The results demonstrate the pressure-dependent nature of perfluorohexane ND activation and confirm that efficient phase transition occurs near the clinical safety limit (MI ≈ 1.9). Scale bar is common to all subfigures.

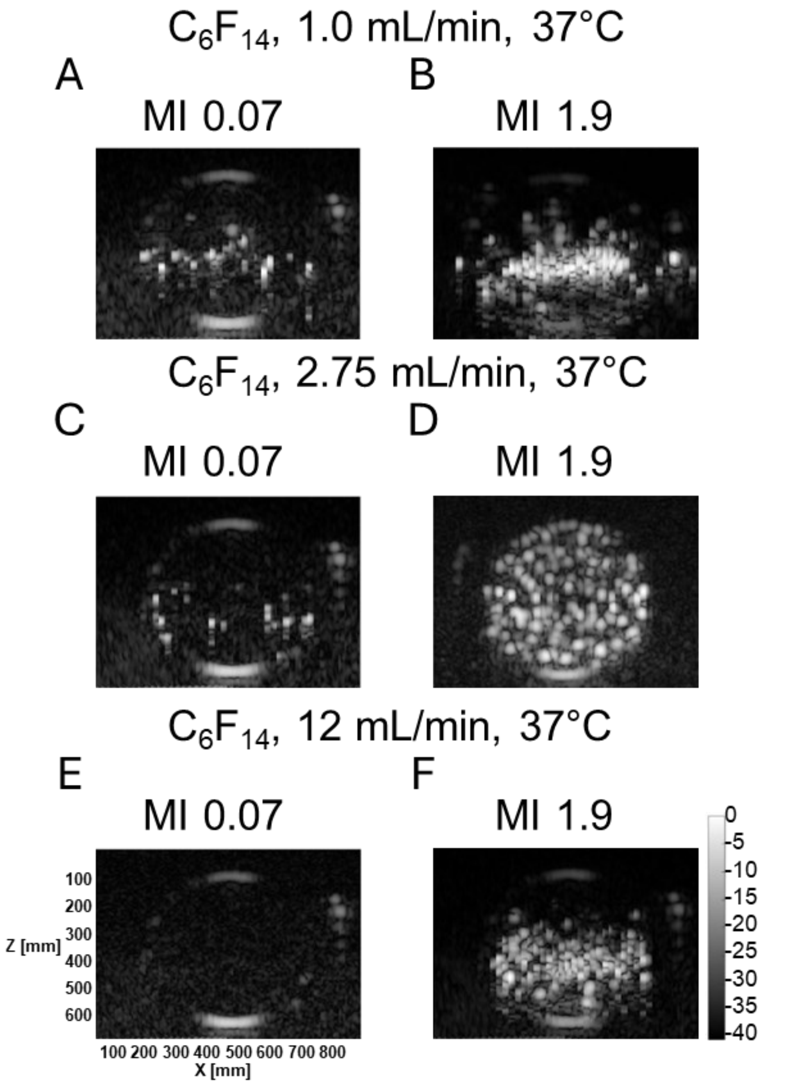


**Figure S4. Effect of microfluidic total flow rate (TFR) on acoustic vaporization behavior of $C_6F_{14}$ nanodroplets at 37 °C.**

Representative B-mode ultrasound images acquired with an L7-4 (5 MHz) linear array transducer showing pressure-dependent acoustic droplet vaporization (ADV) of $C_6F_{14}$ NDs synthesized at different microfluidic total flow rates (TFRs). (A, B) NDs prepared at TFR = 1.0 mL/min. At MI = 0.07, a small degree of spontaneous vaporization and unstable microbubble formation is observed (A). At MI = 1.9, pronounced ADV and dense microbubble clouds form (B). (C, D) NDs prepared at TFR = 2.75 mL/min. Limited spontaneous activation appears at MI = 0.07 but droplets remain more stable than those from the lower-TFR batch (C). Strong ADV is observed at MI = 1.9 (D). (E, F) NDs prepared at TFR = 12 mL/min (NanoAssemblr). At MI = 0.07, no visible vaporization occurs (E), while exposure at MI = 1.9 yields robust ADV (F). Overall, increasing the microfluidic TFR produces smaller, and acoustically stable NDs that require higher acoustic pressure for vaporization, highlighting the influence of synthesis dynamics on ADV thresholds and stability. Scale bar is common to all subfigures.

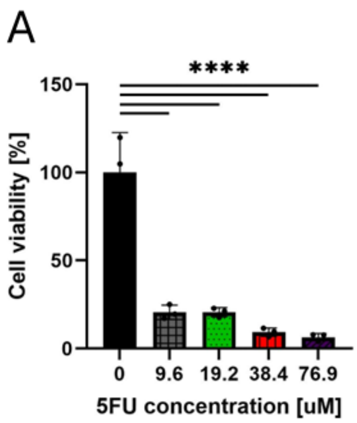


**Figure S5. Cytotoxic effect of free 5-fluorouracil (5-FU) on 4T1 breast cancer cells.**
(A) Cell viability of 4T1 cells after 48-hour exposure to increasing concentrations of free 5-fluorouracil (0 μM, 9.6 μM, 19.2 μM, 38.4 μM, and 76.9 μM). A strong dose-dependent cytotoxic response was observed, with viability decreasing from 100% in the non-treated control to below 20% at 9.6 and 19.2 μM and under 10% at 38.4 and 76.9 μM. These data confirm the potent therapeutic effect of 5-FU against 4T1 tumor cells. Data represent mean ± SD (n = 3). Statistical significance was determined by one-way ANOVA with Tukey's post-hoc test (****$p < 0.0001$).

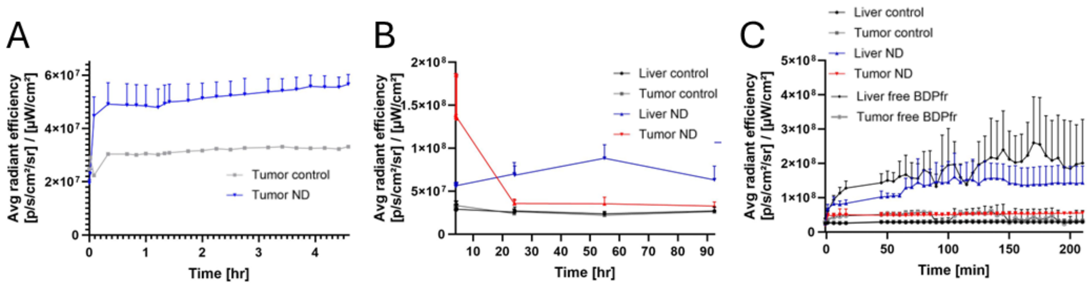


**Figure S6. In vivo fluorescence dynamics and biodistribution kinetics of BDPfr-labeled nanodroplets.**

(A) In vivo IVIS imaging quantification of average radiant efficiency in the tumor region of breast-tumor bearing mice over 4.5 hours post-injection. Mice receiving BDPfr NDs displayed a rapid fluorescence increase within minutes of injection, followed by stable and sustained fluorescence throughout the 4.5 hour imaging period. The signal remained significantly higher than that of the untreated tumor control, indicating rapid accumulation and retention of the NDs in the tumor microenvironment. (B) Long-term in vivo fluorescence kinetics in tumor and liver regions over 90 hours post-injection. Both tumor and liver signals increased sharply immediately following BDPfr ND administration, with the liver exhibiting the highest early-phase accumulation as expected. Fluorescence in the liver decreased to near-baseline (control) levels by 24 hours and remained stable thereafter, suggesting hepatic clearance of free or degraded droplets. In contrast, fluorescence in the tumor continued to rise, showing higher intensity at 24 hours and peaking at approximately 55 hours post-injection before declining slightly by 90 hours, while still remaining above control levels. These results confirm strong and sustained tumor retention of BDPfr NDs compared to transient hepatic uptake. (C) Kinetic profile of average radiant efficiency in liver and tumor tissues over 220 minutes post-injection, illustrating the accumulation and retention dynamics for each experimental group.
